# A conserved ion channel function of STING mediates non-canonical autophagy and cell death

**DOI:** 10.1101/2023.08.26.554976

**Authors:** Jinrui Xun, Zhichao Zhang, Bo Lyu, Defen Lu, Haoxiang Yang, Guijun Shang, Jay Xiaojun Tan

**Author notes:** These authors contributed equally to this work (Jinrui Xun, Zhichao Zhang, Bo Lyu, Defen Lu). Correspondence (G.S.); (J.X.T.).

## Abstract

The cGAS/STING pathway triggers inflammation in response to diverse cellular stresses such as infection, cellular damage, senescence, normal aging, and age-related disease. Besides inflammation, STING also triggers non-canonical autophagy and cell death, the former of which requires the proton pump V-ATPase-mediated LC3 lipidation to single membrane STING vesicles. V-ATPase is known to sense organelle de- acidification in other contexts and recruits the ATG16L1 complex for direct conjugation of LC3/ATG8 to single membranes (CASM). However, it is unclear how STING activates V-ATPase for non-canonical autophagy. Here we report that upon STING activation, the transmembrane domain (TMD) of STING significantly reorganizes and forms an electron-sparse pore in the center. Cellular imaging and in vitro ion flux assays revealed that STING is critical for proton efflux and pH neutralization of Golgi-derived STING vesicles. A chemical ligand of STING, C53, which binds to and blocks the channel of STING strongly inhibited STING-mediated proton flux in vitro and vesicular de-acidification in cells. C53 also abolished STING-dependent LC3 lipidation and cell death. Thus, the ion channel function of STING activates non-canonical autophagy and cell death through vesicle de-acidification.

## Introduction

The cyclic GMP-AMP (cGAMP) synthase (cGAS)/stimulator of interferon genes (STING) innate immunity pathway is critical in host defense against microbial infection^1–4^. Besides infection-triggered inflammation, the cGAS/STING pathway also promotes sterile inflammation in many other conditions involving cytosolic exposure of self-DNA, such as cell damage, senescence, diseases, and normal aging^5,6^. Upon recognition of abnormal cytosolic DNA, cGAS produces a second messenger cGAMP which binds to and activates STING on the endoplasmic reticulum (ER)^7,8^. The cGAMP-bound STING undergoes robust trafficking from the ER to the Golgi region and finally forms perinuclear vesicle clusters where TBK1 is activated to phosphorylate IRF3 and NF-κB, transcription factors that upregulate the expression of type I interferons (IFN-I) and inflammatory cytokines^9^.

STING is also known to restrict microbial infection through non-canonical autophagy, which is independent of autophagy proteins upstream of the ATG5/ATG12/ATG16 complex^10–12^. An extensively studied form of non-canonical autophagy is known as the conjugation of LC3/ATG8 to single membranes or CASM^13^. CASM induction is mediated through the direct recruitment of ATG16 by the vacuolar-type ATPase (V-ATPase), a proton pump that senses organelle de-acidification upon challenges from weak base chemicals, proton ionophores, or lysosomal membrane damage^14–16^. A recent study from the Youle group identified V-ATPase as a downstream effector of STING for non-canonical LC3 lipidation on single membrane STING vesicles^17^. However, it was still unclear how STING activates V-ATPase.

Here, we report that STING-induced non-canonical autophagy depends on an unexpected ion channel function of STING. Ligand-bound STING forms a large electron-poor pore in its transmembrane region that causes proton efflux from post-Golgi STING vesicles and subsequent vesicle de-acidification. This is required for STING-dependent LC3 lipidation and cell death, both of which can be blocked by C53, a small chemical that blocks the transmembrane channel of STING. This new molecular function of STING resembles other CASM-inducing stresses that de-acidify endolysosomes or phagosomes.

## Results

### Activated STING forms a potential channel in its transmembrane region

Our recent structural study of apo-STING and cGAMP/STING complex provided new mechanistic insights into STING autoinhibition and activation^18,19^. Notably, upon cGAMP binding, the ligand binding domain (LBD) of STING contracted, whereas the transmembrane domain (TMD) underwent dilation. After a thorough analysis of the conformational changes in the TMD from the apo and cGAMP-bound structures of human STING, a ligand-induced transmembrane pore with a radius of approximately 1.71 Å at its narrowest part was revealed (**Fig. 1A, B**). In contrast, the pore radius was approximately 0.86 Å at its bottleneck in the apo-structure (**Fig. 1A**). The ligand-induced pore space was electron sparse^18^, suggesting that it is not readily accessible by membrane lipids. Thus, we hypothesized that the TMD of STING might function as an ion channel upon ligand binding.

**Figure. 1.**
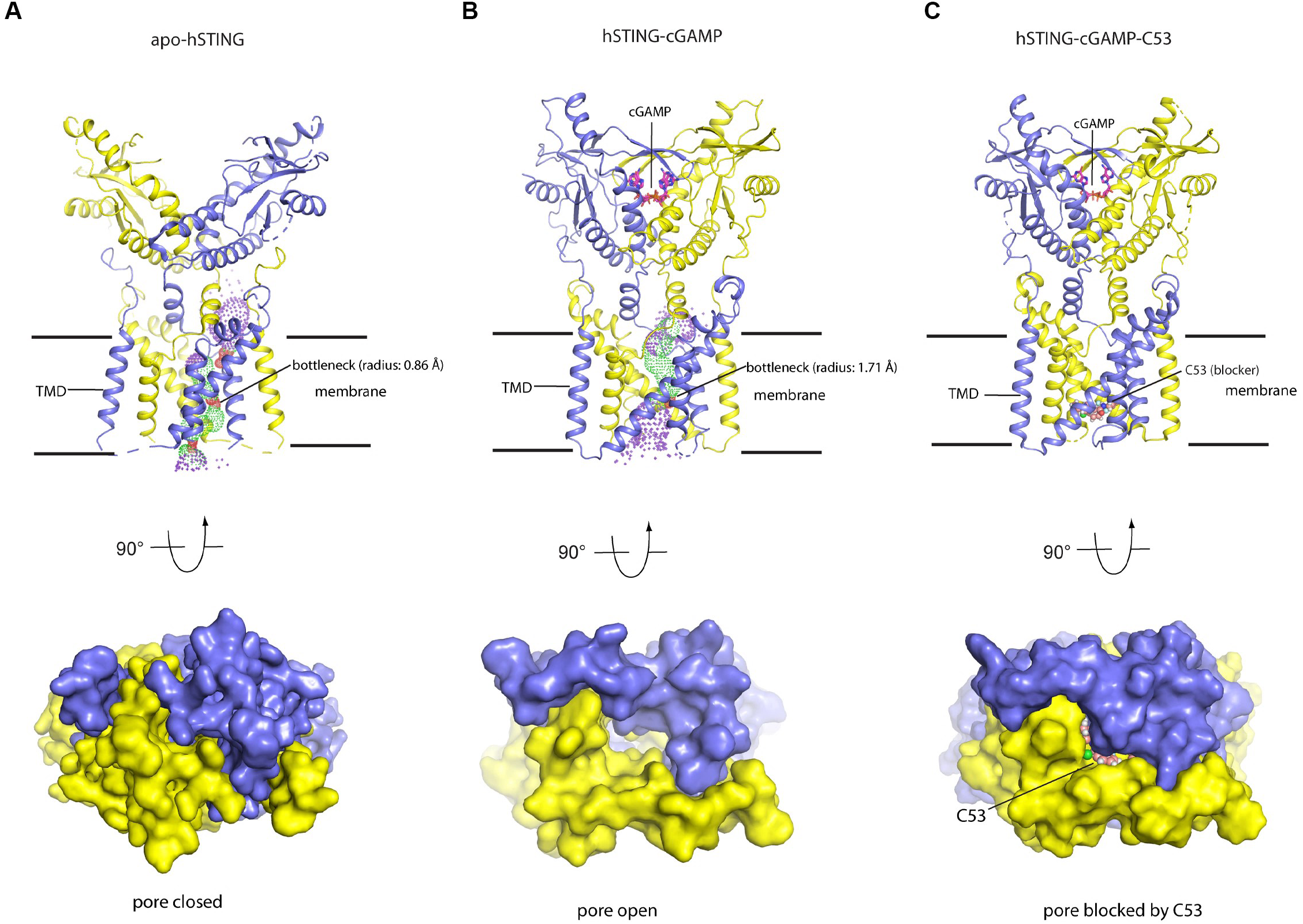
Structural analysis of pore formation in the TMDs of ligand bound and apo-STING. The protomers of STING were shown in cartoon with blue and yellow colors. The cGAMP and C53 were displayed with stick and sphere mode, respectively. The pore radii (spheres) were calculated using HOLE. A. Upper panel, the TMD of human apo-STING (PDB: 6NT5) forms a pore with a radius at its constriction point, its narrowest neck, measuring 0.86 Å; lower panel, the bottom view of pore with surface show. B. Upper panel, the TMD of human cGAMP bound STING (PDB: 8IK3) forms a pore with a radius of 1.71 Å at its constriction point; lower panel, the bottom view of pore with surface show. C. Upper panel, the pore of activated STING (PDB: 7SII) is blocked by compound C53; lower panel, the bottom view of the pore with surface show.

We have previously reported the compound C53 as an unusual STING agonist^20^. C53 binds precisely to the lower part of STING-TMD and appears to fully block the channel (**Fig. 1C**). Thus, we tested C53 as a potential inhibitor of STING’s ion channel function and examined its impact on STING-mediated TBK1 activation as well as LC3 lipidation. Remarkably, the presence of C53 fully abolished STING-dependent LC3 lipidation without affecting TBK1 activation (**Extended Data** Fig. 1A), suggesting that STING-mediated non-canonical autophagy is likely triggered through an ion channel function of STING.

### STING traffics through the Golgi complex upon activation

To explore how the potential STING channel could trigger LC3 lipidation, we first investigated where STING might induce LC3 lipidation in cells. Multiple subcellular localizations have been previously described as active sites for the STING complex, including ER-Golgi-intermediate compartments (ERGIC)^10,21^, the Golgi body^22–24^, and post-Golgi vesicles^17,23,25^. To identify potential organelles where STING might function as an ion channel, we characterized time-dependent STING trafficking upon ligand stimulation. U2OS cells are widely used for immunofluorescence studies, but no STING was expressed in this cell line. We thus established a monoclonal U2OS cell line stably expressing low levels of human STING. This cell line showed robust STING trafficking comparable to endogenous STING in human BJ and HT1080 cells, but is a better model to study STING trafficking as it has consistent STING trafficking throughout the cell population, allowing for careful dissection of the trafficking steps upon ligand stimulation.

Upon cGAMP delivery into U2OS cells, STING showed dramatic subcellular trafficking and strongly colocalized with the cis- and trans-Golgi markers, GM130 and Golgin97, respectively, about 30 min after cGAMP exposure (**Fig. 2A-F**). Remarkably, the occupation of the Golgi membrane by STING caused striking morphological changes of the Golgi complex, as shown by more extensive overlap between GM130 and Golgin97 (**Extended Data** Fig. 1B-D) as well as swelled Golgi stacks at 30 min (**Fig. 2A, D, Extended Data** Fig. 1B, E). After this time, a large number of STING vesicles formed out of the Golgi complex (**Fig. 2A-F**), and the Golgi markers returned back to resting morphology (**Extended Data** Fig. 1B-E). Despite the extensive clustering of Golgi-derived STING vesicles that appeared to partially overlap with Golgi markers, these STING vesicles are apparently different membranes outside of the Golgi stacks (**Fig. 2A, D**).

**Figure 2.**
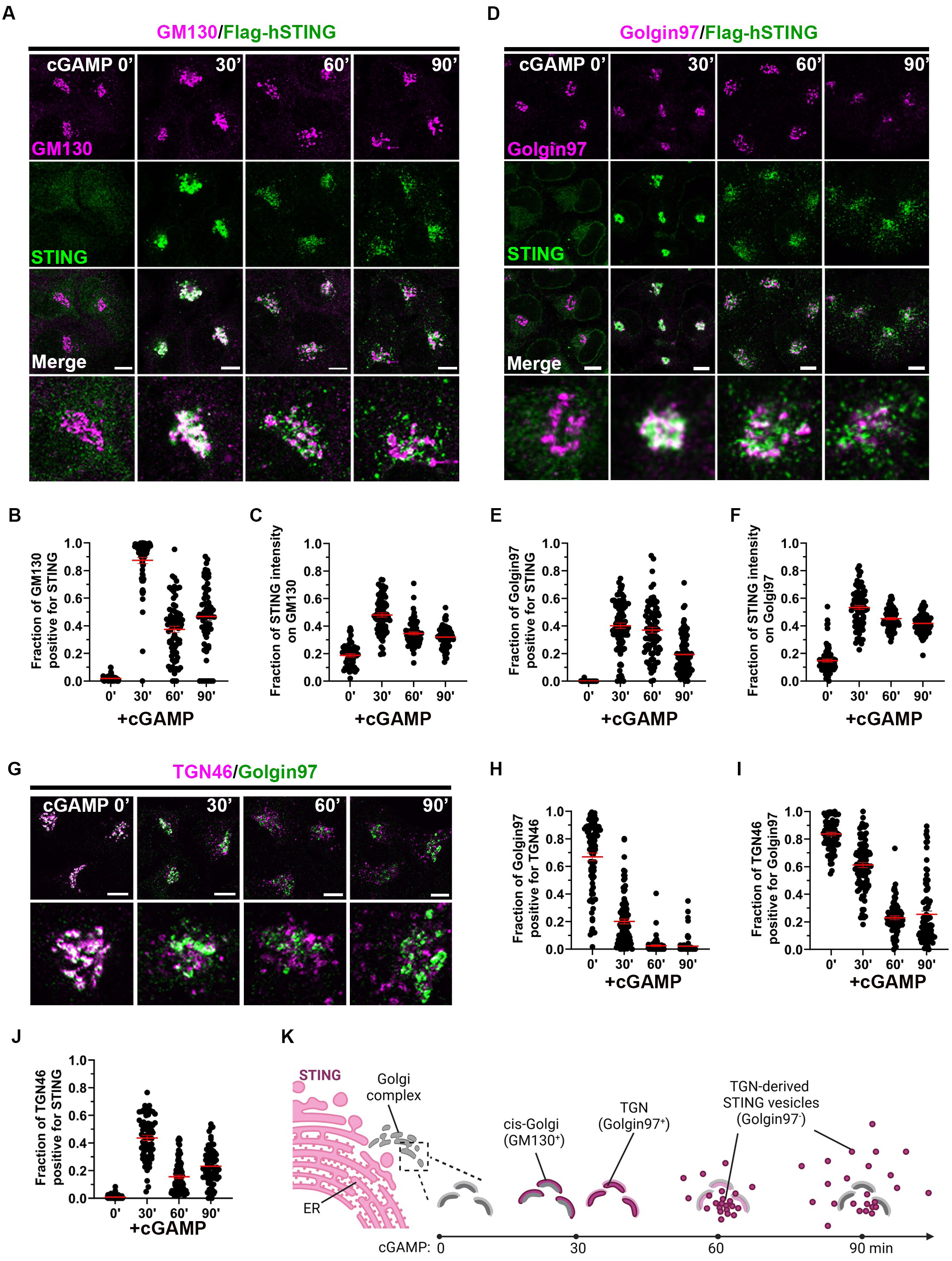
Activated STING traffics through the Golgi complex. **A.** STING traffics through the *cis*-Golgi upon cGAMP binding. Monoclonal U2OS cells stably expressing Flag-tagged human STING (hSTING) were stimulated with 1 μM cGAMP and fixed at indicated time points for the co-staining of STING and the cis-Golgi marker GM130. **B, C.** Quantification of the colocalization between STING and GM130 in (A). Mean ± SEM; n = 53, 80, 79, and 67 random cells for 0’, 30’, 60’, and 90’, respectively. **D.** STING traffics through the *trans*-Golgi upon cGAMP binding. U2OS Flag-hSTING cells the same as in (A) were stimulated with 1 μM cGAMP and fixed at indicated time points for the co-staining of STING and the trans- Golgi marker Golgin97. **E, F.** Quantification of the colocalization between STING and Golgin97 in (D). Mean ± SEM; n = 81, 89, 82, and 88 random cells for 0’, 30’, 60’, and 90’, respectively. **G.** STING trafficking triggers TGN46 budding from the *trans*-Golgi. U2OS Flag-hSTING cells the same as in (A) were stimulated with 1 μM cGAMP and fixed at indicated time points for the co-staining of two different TGN markers TGN46 and Golgin97. **H, I.** Quantification of the colocalization between TGN46 and Golgin97 in (G). Mean ± SEM; n = 83, 100, 77, and 90 random cells for 0’, 30’, 60’, and 90’, respectively. **J.** Quantification of the colocalization between TGN46 and STING in U2OS Flag-hSTING cells. Mean ± SEM; n = 80, 66, 80, and 70 random cells for 0’, 30’, 60’, and 90’, respectively.**K.** Schematic illustration of STING trafficking through the Golgi stacks to post-Golgi vesicles. Created using Biorender. Bar, 10 μm.

STING trafficking caused robust relocalization of the trans-Golgi network (TGN) marker TGN46 (also known as TGN38 or Trans-Golgi Network Protein 2, TGOLN2). While TGN46 showed extensive overlap with Golgin97 in resting conditions, the arrival of STING to the Golgi 30 min after cGAMP binding is accompanied by the budding of TGN46, but not Golgin97, from the trans-Golgi stack (**Fig. 2G-I**). The TGN46 vesicles and Golgi-derived STING vesicles appeared to be distinct but with partial association with each other (**Fig. 2J and Extended Data Fig. S1F**).

STING trafficking through the Golgi complex was also observed when hSTING was activated by diABZI (**Extended Data** Fig. 1G, H). Similarly, endogenous STING in HT1080 and BJ cells were both found to traffic through the Golgi upon cGAMP treatment (**Extended Data** Fig. 2A, B). Mouse STING (mSTING) stably expressed in U2OS also trafficked through the Golgi upon cGAMP activation (**Extended Data** Fig. 2C-H). Thus, in different cell lines, ligand binding activates robust STING trafficking through the Golgi complex to Golgi-derived vesicles (**Fig. 2K**).

### STING-induced LC3 puncta mostly localize to post-Golgi, endosome-like vesicles

Since STING showed robust trafficking through the Golgi stacks, we asked whether STING stimulated LC3 lipidation directly on the Golgi membrane. Surprisingly, cGAMP-stimulated endogenous LC3 puncta had little overlap with the Golgi (**Fig. 3A-C, Extended Data** Fig. 3A-C). The LC3 puncta did not appear 30 min after cGAMP exposure when most STING accumulated at the Golgi (**Fig. 3A, Extended Data** Fig. 3A, C), and their subsequent occurrence correlated with STING trafficking to post-Golgi vesicles (**Fig. 2A-F**). The morphology of the triggered LC3 puncta resembled post-Golgi clusters of STING vesicles (**Fig. 3A**, **Fig. 2A, D, Extended Data** Fig. 3A, C). Indeed, three channel imaging showed that these LC3 puncta largely colocalized with STING but not Golgin97 (**Extended Data** Fig. 3D, E). Thus, these data strongly support that STING-stimulated LC3 lipidation primarily on post-Golgi vesicles.

**Figure 3.**
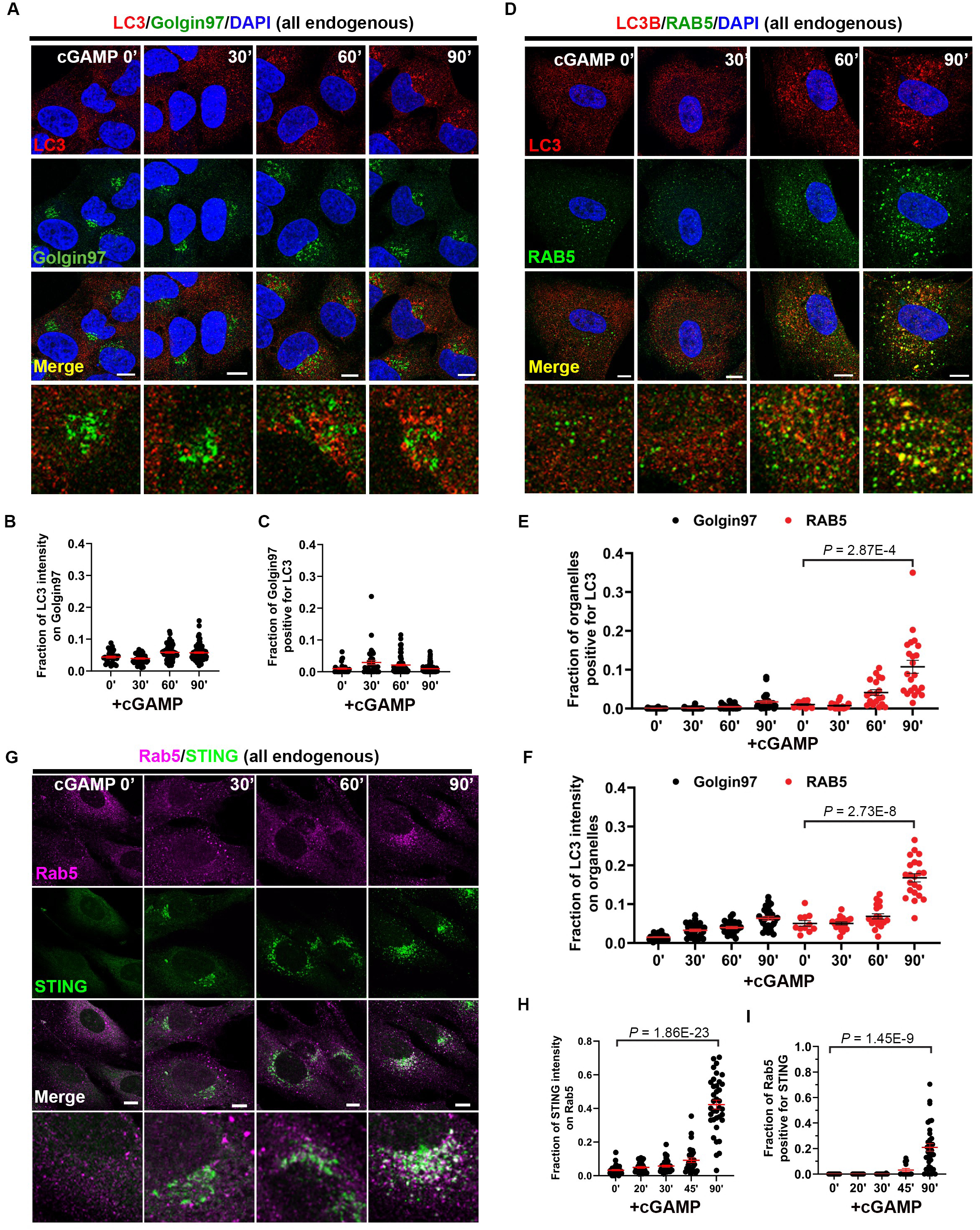
STING induces LC3 puncta on post-Golgi, endosome-like vesicles. **A.** STING induces LC3 puncta outside of the Golgi. Monoclonal U2OS Flag-hSTING cells were stimulated with 1 μM cGAMP and fixed at indicated time points for the co-staining of endogenous LC3B and the *trans*-Golgi marker Golgin97. **B, C.** Quantification of the colocalization between LC3 and Golgin97 in (A). Mean ± SEM; n = 31, 40, 72, and 92 random cells for 0’, 30’, 60’, and 90’, respectively. **D.** STING-induced LC3 puncta colocalize with RAB5. BJ cells with endogenous STING were stimulated with 1 μM cGAMP and fixed at indicated time points for the co-staining of endogenous LC3 and the early endosome marker RAB5. **E, F.** Quantification of the colocalization between LC3B and RAB5 in (D) or Golgin97 in Extended Data Fig. 3C. Mean ± SEM; n = 31, 30, 25, 27, 11, 18, 19, and 22 random cells from left to right. **G.** Post-Golgi STING puncta colocalize with RAB5. BJ cells were stimulated with 1 μM cGAMP and fixed at indicated time points for the co-staining of endogenous STING and RAB5. **H, I.** Quantification of the colocalization between STING and RAB5 in (G). Mean ± SEM; n = 39, 27, 35, 31, 37, 35, 15, 21, 13, and 34 random cells from left to right. Bar, 10 μm.

These observations are consistent with recent findings from the Youle group that cGAMP stimulates LC3 lipidation on single membrane STING vesicles^17^. To further explore the identity of these vesicles, we co-stained LC3 with different organelle markers at different time points after cGAMP treatment. This revealed a significant colocalization between the triggered LC3 puncta with the early endosome marker RAB5, but not Golgin97 (**Fig. 3D-F, Extended Data** Fig. 3C). Of note, the total punctate intensities of RAB5 substantially increased after STING leaves the Golgi (**Fig. 3D**), suggesting that post-Golgi STING vesicles likely developed endosome-like properties and might recruit RAB5. Consistently, a sharp increase in STING colocalization with RAB5 was also observed in its post-Golgi stage (**Fig. 3G-I**). Further imaging analysis revealed that cGAMP-induced LC3 puncta were typically found on STING-vesicles positive for RAB5 (**Extended Data** Fig. 3F). STING also developed increased colocalization with the late endosome/lysosome markers CD63 and LAMP1, but to a much lower extent compared with RAB5 (**Extended Data** Fig. 3G-L), which might reflect a partial late endosome properties of STING vesicles or an indication of STING delivery to the lysosome for degradation. Together, these results suggest a gradual acquisition of endolysosomal properties by post-Golgi STING vesicles where most LC3 puncta are detected.

### STING de-acidifies post-Golgi vesicles

Our data suggest that STING triggered LC3 lipidation mostly on post-Golgi, endosome-like vesicles. Given that C53, a chemical that binds to the transmembrane pore of STING, selectively blocks STING-mediated LC3 lipidation without affecting TBK1 activation or subsequent STING phosphorylation (**Extended Data** Fig. 1A), we hypothesized that a potential ion channel function of STING triggers LC3 lipidation on post-Golgi STING vesicles. STING-mediated LC3 lipidation is known to depend on the V-ATPase proton pump that directly recruits ATG16L1^17^. In other contexts, the same V-ATPase-ATG16L1 axis senses the de-acidification of peripheral organelles such as phagosomes and endolysosomes and triggers direct LC3 lipidation onto these compartments, known as CASM^13,16^. Since STING also activates CASM through V-ATPase, we examined whether STING activation could raise the pH of its post-Golgi trafficking vesicles.

Lyso-pHluorin is a pH sensor of endolysosomes, the fluorescence of which is quenched at acidic pH and activated upon pH neutralization or endolysosomal de-acidification^26^. We tested if lyso-pHluorin could serve as a pH sensor of STING vesicles since they developed endolysosomal properties (**Fig. 3**). Remarkably, STING activation triggered the formation of lyso-pHluorin puncta in both U2OS cells reconstituted with human STING (hSTING) and BJ cells expressing endogenous hSTING (**Fig. 4A, B and Extended Data** Fig. 4A). Similar lyso- pHluorin puncta were observed when mouse STING was expressed in U2OS cells and activated by DMXAA, a chemical agonist selectively activating mouse STING^27–29^ (**Fig. 4C, D**). The activation of neither human nor mouse STING caused any puncta formation of EGFP-galectin3 (**Extended Data** Fig. 4B), an endolysosomal membrane damage sensor, suggesting that the membrane integrity of STING vesicles was compromised and that the observed pH neutralization was likely due to the altered activity of potential ion channels. The vesicle de-acidification activity of STING appeared to be highly conserved, given that lyso-pHluorin puncta were equally detected when *Xenopus tropicalis* STING (XtSTING) was activated by cGAMP in U2OS cells (**Fig. 4E, F**). We consistently noticed the development of brighter, STING-dependent lyso-pHluorin puncta over time. This was confirmed using time lapse live cell imaging that showed substantial lyso-pHluorin puncta 60-120 min after the activation of mSTING (**Fig. 4G, H**), consistent with STING trafficking to endosome-like vesicles at this time (**Fig. 3**).

**Figure 4.**
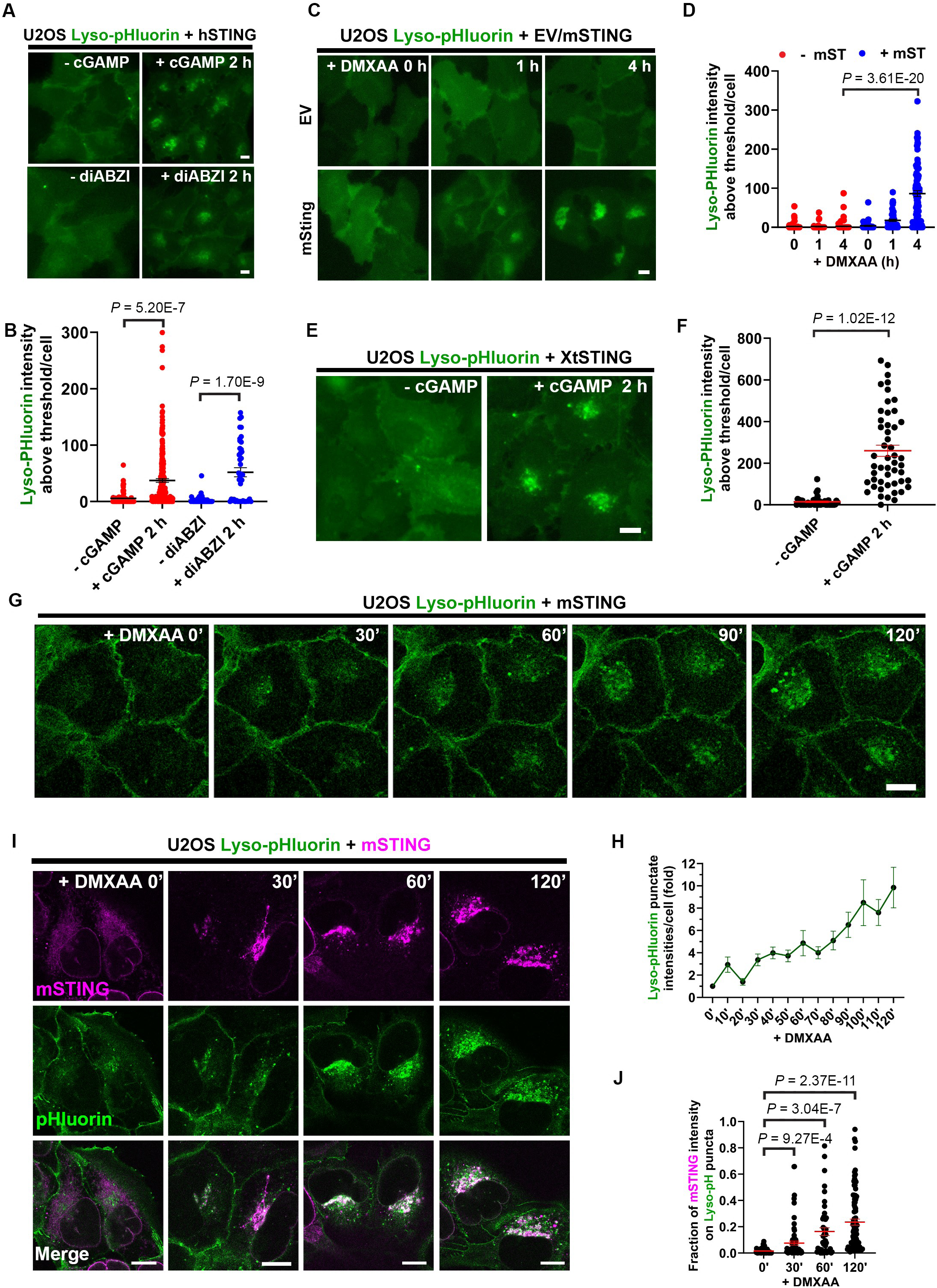
STING de-acidifies its post-Golgi trafficking vesicles. **A.** Activation of human STING (hSTING) triggers lyso-pHluorin puncta. U2OS cells stably expressing hSTING and a genetically encoded pH sensor lyso-pHluorin were stimulated as indicated and the lyso-pHluorin puncta were monitored by live cell imaging. **B.** Quantification of the average intensities of lyso-pHluorin puncta in (A). Mean ± SEM; n = 73, 249, 50, and 42 random cells from left to right. **C.** Activation of mouse STING (mSTING) triggers lyso-pHluorin puncta. U2OS cells stably expressing mSTING and lyso-pHluorin were stimulated with DMXAA and the lyso-pHluorin puncta were monitored by live cell imaging. **D.** Quantification of the average intensities of lyso-pHluorin puncta in (C). Mean ± SEM; n = 70, 44, 83, 44, 53, 89 random cells from left to right. **E.** Activation of Xenopus tropicalis STING (XtSTING) triggers lyso-pHluorin puncta. U2OS cells stably expressing XtSTING and lyso-pHluorin were stimulated with cGAMP and the lyso-pHluorin puncta were monitored by live cell imaging. **F.** Quantification of the average intensities of lyso-pHluorin puncta in (E). Mean ± SEM; n = 50 (- cGAMP), 42 (+ cGAMP 2 h) random cells. **G.** Time lapse confocal live cell imaging shows mSTING-induced lyso-pHluorin puncta upon DMXAA treatment. **H.** Quantification of the average intensities of lyso-pHluorin puncta in (G). Mean ± SEM; n = 32 cells. **I.** STING-induced lyso-pHluorin puncta colocalize with STING. U2OS cells stably expressing Flag-mSTING and lyso-pHluorin were stimulated with DMXAA and then fixed at indicated time points for the staining of mSTING by Flag antibody. See also Extended Data Fig. S4C. **J.** Quantification of the colocalization between STING and lyso-pHluorin in (I). Mean ± SEM; n = 55, 76, 47, and 93 random cells from left to right. Bar, 10 μm.

To further characterize whether the observed lyso-pHluorin puncta represent STING vesicles or other nearby, pre-existing endolysosomes, we developed a protocol to stain for STING in the same cells after the induction of lyso-pHluorin puncta by cGAMP. We fixed the cells so that the pre-existing fluorescence of lyso-pHluorin was not changed by the fixation and immunostaining steps (**Extended Data** Fig. 4C). This approach revealed extensive colocalization between cGAMP-induced lyso-pHluorin puncta and STING (**Fig. 4H, I**). Thus, post-Golgi STING vesicles are de-acidified by STING, which might be sensed by the V-ATPase proton pump for direct LC3 lipidation on these vesicles. Consistent with vesicle de-acidification being upstream of LC3 lipidation, STING-induced lyso-pHluorin puncta formation was normal in ATG5- and ATG7-knockout cells (**Extended Data** Fig. 4D, E).

### STING mediates proton flux in vitro

The vesicle de-acidification activity of STING further suggests that the transmembrane pore of STING might function as an ion channel allowing protons and/or other ions to pass through the membrane, which is consistent with our structural analysis (**Fig. 1**). To test this hypothesis, we conducted an in vitro ion flux assay^30^ using purified STING reconstituted onto proteoliposomes (**Fig. 5A**). In this assay, proton influx was driven by a strong gradient for the efflux of potassium across the membrane. The proton sensitive dye 9-Amino-6-chloro-2- methoxyacridine (ACMA) was used to monitor the pH change inside the vesicles. A potassium ionophore valinomycin was added to initiate the flux. Surprisingly, in this system, STING showed constitutive activity in proton flux, which was not further enhanced by the addition of diABZI, a membrane-permeable STING agonist expected to activate STING from the luminal side of the liposomes (**Fig. 5B**). Nevertheless, the basal proton flux activity was strongly inhibited by C53 (**Fig. 5B**) which binds to and blocks the transmembrane pore of STING (**Fig. 1C**). Taken together, our structure analysis, cellular imaging, and in vitro ion flux assay suggest that STING functions as a proton channel to de-acidify at least Golgi-derived vesicles, consistent with cGAMP-induced STING trafficking to perinuclear acidic compartments.

**Figure. 5.**
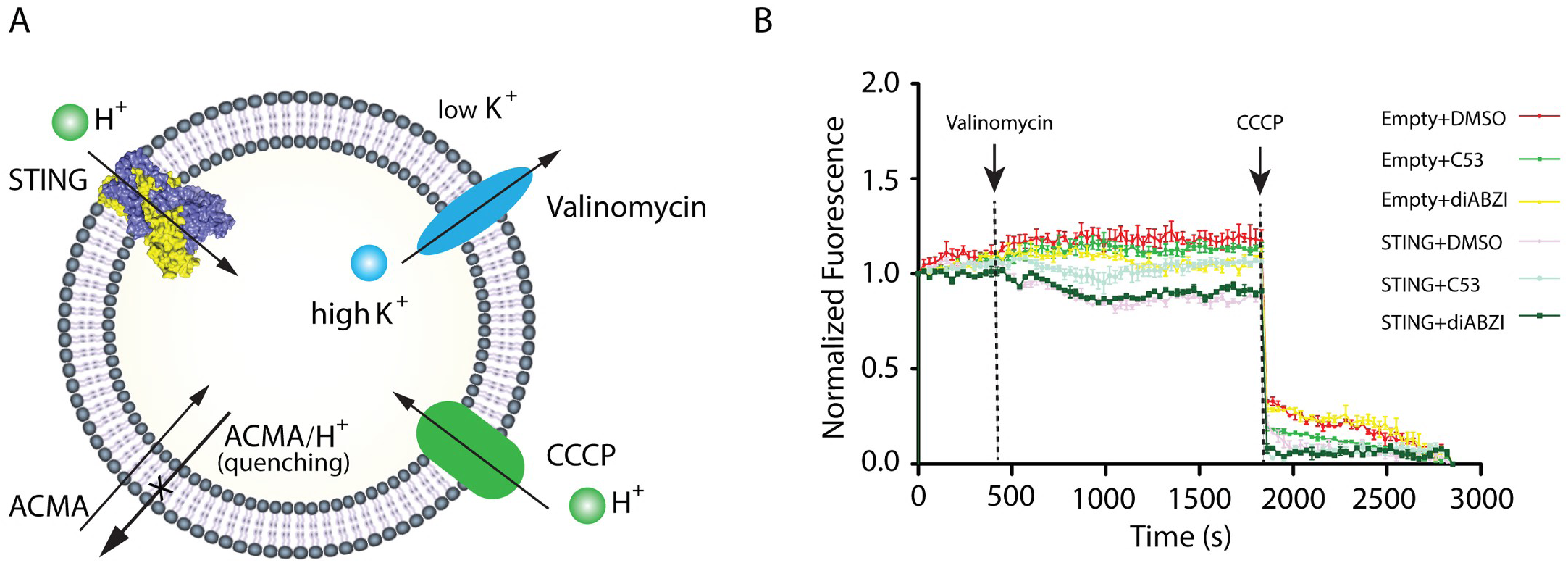
In vitro analysis of human STING as a proton channel. **A.** Schematic illustration of the fluorescence-based proton flux assay driven by a K^+^ gradient. Vesicles were loaded with 450 mM KCl and then diluted into flux buffer with 450 mM NaCl in the presence of ACMA. Valinomycin was added to initiate the flux and proton-ionophore carbonyl cyanide chlorophenylhydrazone (CCCP) was used to collapse the electrical gradient. **B.** Human STING-containing vesicles (1:100 wt/wt protein-to-lipid ratios) were diluted into buffer containing 450 mM NaCl. Valinomycin and CCP were added at ∼300 s and 1800 s, respectively (arrowhead and grey dashed line) and fluorescence was detected over time. Empty (without STING protein) vesicles were used as control. Data are shown as mean ± s.e.m. n = 6 independent experiments.

STING-mediated ion flux is likely selective for protons, unique ions with an extremely small size. First, the heavily positively charged cytosolic side of this channel would not allow the efflux of cation ions such as Na+ and K+ (**Extended Data** Fig. 5A). Second, the relatively small size of the pore might preclude the passage of anion ions like hydrated Cl- with an estimated radius of 3.32 Å. Finally, mutations of various amino acid residues in the center of the transmembrane pore in human STING (**Extended Data** Fig. 5B, C) failed to identify any mutants completely dead in vesicle de-acidification (**Extended Data** Fig. 5D, E), among which those triggered weaker or no lyso-pHluorin puncta turned out to be unstable (**Extended Data** Fig. 5F). One mutant L54E induced brighter lyso-pHluorin puncta than wild type and any other mutants of STING (**Extended Data** Fig. 5C), suggesting increased proton efflux by this mutant. As the L54E mutation is located near the narrowest neck of the channel (**Extended Data** Fig. 5G), the increased proton efflux is likely mediated by an interaction between protons and E54 that allows more efficient proton passage through the channel.

### The channel of STING mediates vesicle de-acidification and LC3 lipidation

The data strongly argue for an ion channel function of STING that de-acidifies its post-Golgi trafficking vesicles, thus allowing for the activation of the V-ATPase-ATG16L1 axis for LC3 lipidation. Since we were not able to find a STING mutant with a selective loss-of-function in vesicle de-acidification (**Extended Data** Fig. 5C-E), we next focused on the STING channel blocker C53 in characterizing the role of ion flux in vesicle de-acidification and LC3 lipidation.

For C53 to be a valid blocker of the ion channel function of STING, we first need to rule out its potential impact on STING trafficking from the ER to post-Golgi vesicles. We have recently shown that although C53 binds to the transmembrane pore of human STING, it is also a STING agonist that triggers STING trafficking from the ER, leading to TBK1 activation^20^. We thus examined whether STING trafficking was different with or without C53. Surprisingly, C53 alone induced large numbers of STING puncta with little colocalization with the Golgi (**Extended Data** Fig. 6A). The addition of C53 together with either cGAMP or diABZI prevented STING trafficking to the Golgi, causing the accumulation of both endogenous and ectopic STING in punctate structures similarly to what was stimulated by C53 alone (**Fig. 6A-C, Extended Data** Fig. 6A-D). Thus, the presence of C53 redirects STING to an abnormal route that blocks STING trafficking at a pre-Golgi stage.

**Figure 6.**
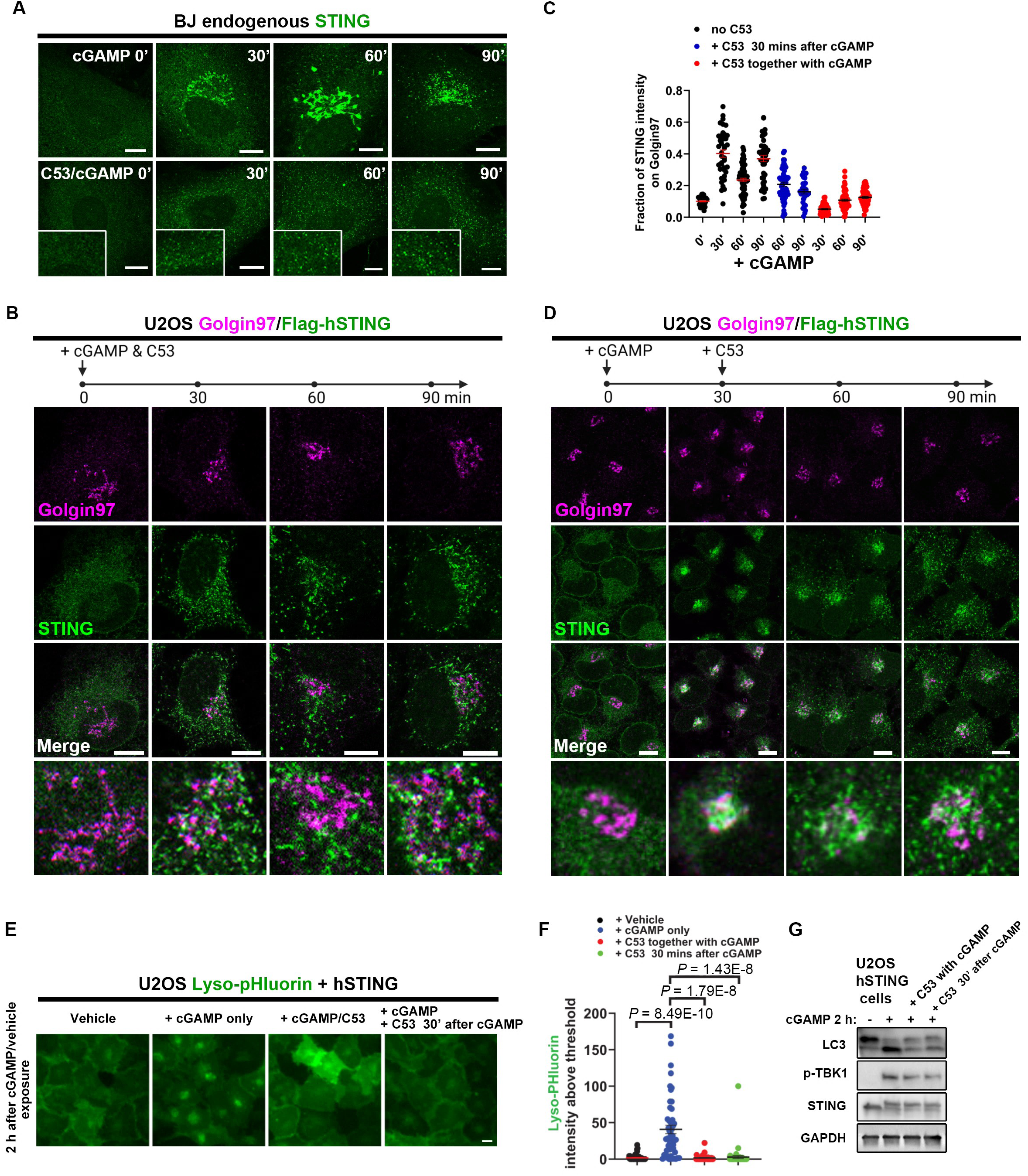
The STING channel is required for vesicle de-acidification and LC3 lipiadtion. **A.** Compound C53, which binds to the transmembrane pore of STING, fully blocks the trafficking of endogenous STING from the ER to the Golgi in BJ cells. BJ cells were stimulated with cGAMP alone or cGAMP + C53 for indicated time periods, followed by fixation and immunostaining of STING. **B.** Compound C53 blocks Flag-hSTING trafficking to the Golgi. Monoclonal U2OS Flag-hSTING cells were stimulated with cGAMP together with C53 for indicated time periods and then fixed for the immunostaining of STING and Golgin97. **C.** Quantification of the colocalization between STING and Golgin97 in different conditions. Mean ± SEM; n = 28, 39, 62, 44, 56, 32, 50, 64, and 56 random cells from left to right. Note that C53 addition 30 min after cGAMP does not block STING trafficking to or from the Golgi, but adding C53 together with cGAMP blocked STING trafficking to the Golgi. **D.** C53 addition 30 min after cGAMP allows STING trafficking to the Golgi and post-Golgi vesicles. Monoclonal U2OS cells stably expressing hSTING were stimulated as indicated and then fixed for the immunostaining of STING and Golgin97. **E.** C53 addition 30 min after cGAMP fully blocks STING-induced lyso-pHluorin puncta. U2OS cells stably expressing hSTING and lyso-pHluorin were stimulated as indicated and the lyso-pHluorin puncta were monitored by live cell imaging. **F.** Quantification of the average intensities of lyso-pHluorin puncta in (E). Mean ± SEM; n = 63, 57, 55, and 76 random cells from left to right. **G.** C53 addition 30 min after cGAMP fully blocks STING-induced LC3 lipidation. Monoclonal U2OS Flag-hSTING cells were stimulated with cGAMP and C53 as indicated and all cells were harvested at 2 hours for western blot analysis of indicated proteins. Bar, 10 μm.

To assess the impact of C53 without blocking STING trafficking, we tested C53 addition after STING arrived at the Golgi. In U2OS Flag-STING monoclonal cells, most STING accumulated at the Golgi 30 min after cGAMP treatment (**Fig. 2, 6C, D**). Adding C53 at this time point did not block STING trafficking to post-Golgi vesicles (**Fig. 6C, D, Extended Data** Fig. 6C, E). However, C53 still almost completely blocked STING-dependent vesicle de-acidification (**Fig. 6E, F**) as well as LC3 lipidation, without affecting TBK1 activation (**Fig. 6G**). Compared with cGAMP, diABZI triggered slower STING trafficking, with most STING found at the Golgi 60 min after stimulation (**Extended Data** Fig. 1G, H). Adding C53 at this time still fully blocked diABZI-induced LC3 lipidation (**Extended Data** Fig. 6F).

We further tested adding C53 after STING fully arrived at post-Golgi vesicles. Specifically, we added C53 two hours after cGAMP or diABZI when vesicle de-acidification was already detected. Remarkably, C53 still dramatically suppressed STING-mediated vesicle de-acidification, and even fully cleared diABZI-induced lyso-pHluorin puncta (**Fig. 7A-B**). Under these conditions, STING-mediated LC3 lipidation was not reversed by C53 as it was already saturated by 2 hours (**Extended Data** Fig. 6G). Taken together, after STING traffics to the Golgi or post-Golgi vesicles, its channel blocker C53 selectively suppresses STING-mediated vesicle de- acidification and LC3 lipidation, without affecting TBK1 signaling.

**Figure 7.**
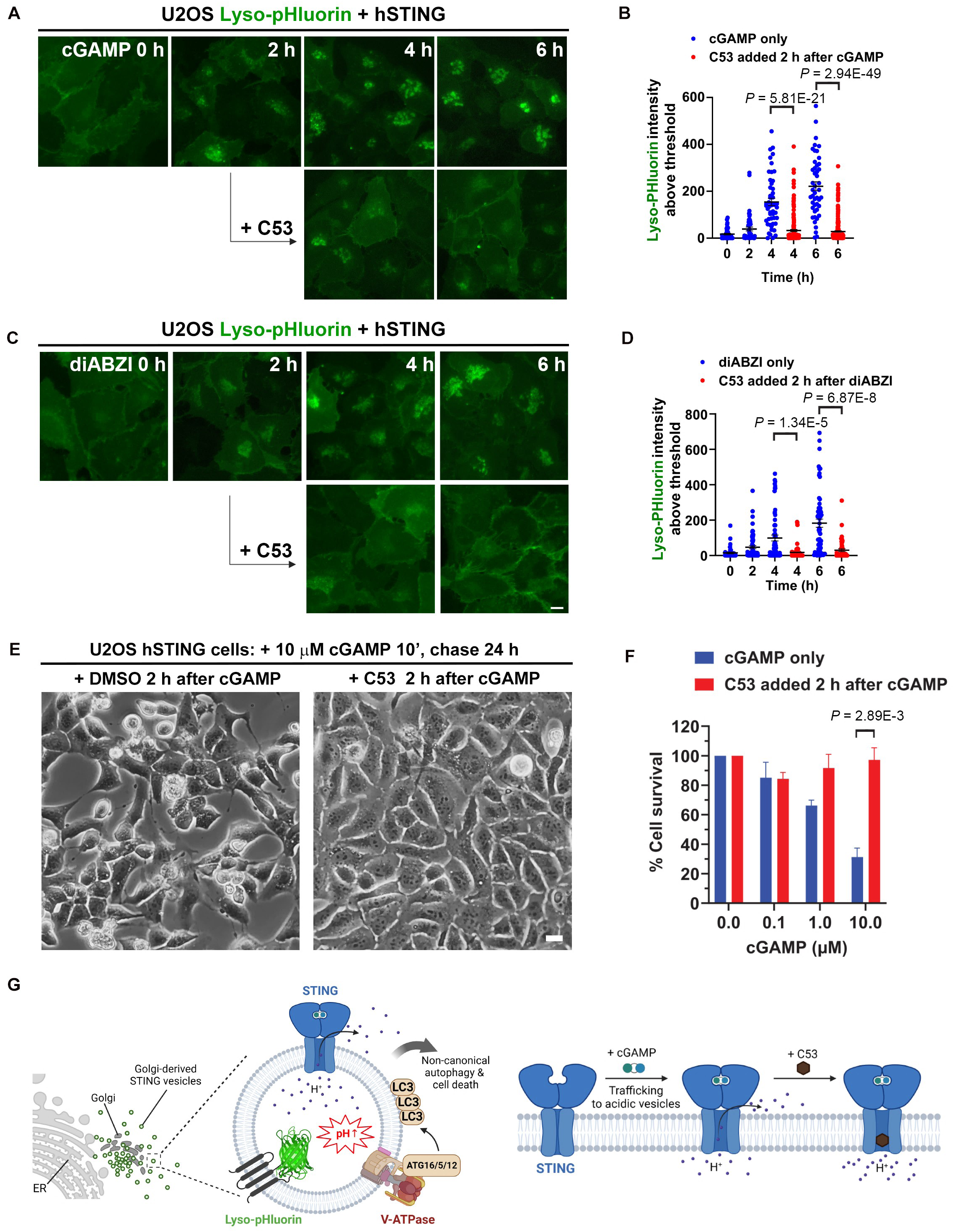
Blocking the STING channel reverses vesicle de-acidification and prevents STING-mediated cell death. **A.** C53 addition 2 hours after cGAMP treatment strongly suppresses STING-induced lyso-pHluorin puncta. U2OS cells stably expressing hSTING and lyso-pHluorin were stimulated as indicated and the lyso-pHluorin puncta were monitored by live cell imaging. **B.** Quantification of the average intensities of lyso-pHluorin puncta in (A). Mean ± SEM; n = 50, 51, 51, 191, 46, and 257 random cells from left to right. **C.** C53 addition 2 hours after diABZI treatment strongly suppresses and reverses STING-induced lyso-pHluorin puncta. Cells were treated similarly as in (A) except that STING was activated by diABZI. **D.** Quantification of the average intensities of lyso-pHluorin puncta in (C). Mean ± SEM; n = 42, 62, 65, 61, 61, and 50 random cells from left to right. **E.** C53 addition 2 hours after cGAMP treatment fully blocks STING-dependent cell death. Monoclonal U2OS Flag-hSTING cells were treated as indicated and the cell death was analyzed 24 hours after cGAMP treatment by cell imaging. **F.** Quantification of the cell death assays in (E). Cells were fixed, permeabilized, and stained with DAPI. The average number of cells in each microscope field of view was automatically counted by software. Percentages of cells compared with no cGAMP control were presented. Mean ± SEM. n = 3 independent experiments. **G.** Schematic model of the ion channel function of STING in vesicle de-acidification, non-canonical autophagy, and cell death. Left, activated STING traffics through the Golgi complex to perinuclear clusters of endosome-like acidic vesicles; the channel function of STING causes proton efflux and vesicle de-acidification, which can be sensed by a pH sensor lyso-pHluorin; the increased luminal pH of STING vesicles activates the V-ATPase- ATG16L1 axis for non-canonical LC3 lipidation onto STING vesicles; activation of the STING channel also triggers cell death. Right, cGAMP induces the enlargement of the transmembrane pore of STING as well as STING trafficking to post-Golgi acidic vesicles; compound C53 can be used to block the STING channel when added after STING trafficking to the Golgi or post-Golgi vesicles. Created using Biorender. Bar, 10 μm.

### STING-mediated cell death depends on its transmembrane channel

Over activation of STING is known to cause cell death through an interferon-independent function of STING^31^. A C-terminal “RHLR” motif of STING required for its trafficking from the ER is also critical for STING-dependent cell death^10,31^. We recapitulated STING-mediated cell death in U2OS cells stably expressing hSTING and examined whether it required the ion channel function of STING (**Fig. 7E**). Cells were briefly treated with cGAMP/digitonin buffer for 10 min and then changed back to original media. Extensive cell death was observed 24 hours after cGAMP treatment (**Fig. 7E, F**). Strikingly, the addition of C53 two hours after cGAMP treatment fully blocked cGAMP-induced cell death (**Fig. 7E, F**). It is of note that C53 alone, which is sufficient to activate TBK1 signaling for interferon production, did not cause any cell death in these tests (**Fig. 7E, F**). Thus, STING-mediated cell death is dependent on its ion channel function but independent of TBK1 signaling.

In summary, we discovered an unexpected ion channel function of STING that allows the efflux of protons from post-Golgi STING vesicles, leading to vesicle de-acidification, non-canonical autophagy, and cell death, all independent of STING-induced TBK1/interferon signaling (**Fig. 7G**). STING-mediated vesicle de-acidification can be captured by lyso-pHluorin, a genetically-encoded lysosomal pH sensor that is also targeted to post-Golgi STING vesicles. Although the channel blocker C53 disrupts STING trafficking, its addition after STING arrives at the Golgi or post-Golgi vesicles can selectively block the ion channel functions of STING. The new function of STING appears to be highly conserved as STING from humans, mice, and frogs all stimulated vesicle de- acidification when activated (**Fig. 4A-F**). Such changes in vesicle pH likely activate the V-ATPase-ATG16L1 axis for LC3 lipidation onto STING vesicles^17^, resembling other CASM inducers^14–16^.

## Discussion

How STING triggers non-canonical autophagy and cell death are fundamental questions in innate immunity. In this study, through structural biological, cell biological, and biochemical approaches, we found that activated STING forms a transmembrane channel that mediates both non-canonical autophagy and cell death. The channel of STING allows efflux of protons, leading to de-acidification of post-Golgi STING vesicles. Vesicle de- acidification is a well-established stimulus that activates the assembly of the V-ATPase proton pump which directly recruits ATG16L1 for ATG8 lipidation onto the local membrane^13–16^. Consistent with our findings, the V-ATPase-ATG16L1 axis has been previously reported to mediate STING-dependent LC3 lipidation^17^. Thus, STING-mediated vesicle de-acidification mimics other cellular stresses such as lysosomal membrane damage or exposure to proton ionophores, instigating the same downstream effectors for the induction of non-canonical autophagy or CASM^13–16^.

Our study suggests that the proton channel function of STING is highly conserved. A recent study identified a bacterial cGAMP receptor as a potential cGAMP-activated Cl^-^ channel^32^. The surface charges on the cytosolic side of the pore suggest that STING might also allow for the flux of anions like Cl^-^ in addition to H^+^. Alternatively, anion-binding to the cytosolic side of the pore might allow for more efficient H^+^ leakage. In addition, the size of the STING channel might change when it is transported to acidic post-Golgi vesicles, which in turn would affect the channel selectivity in ion flux. Thus, future studies are needed to thoroughly test the ion selectivity of the STING channel.

The extensive clustering of Golgi-derived STING vesicles near the microtubule-organization center has rendered it difficult to differentiate post-Golgi STING vesicles from the Golgi stacks themselves in lower resolution microscopy. However, our in-depth analysis of ligand-stimulated STING trafficking in different cell lines strongly suggests the active sites of STING-mediated LC3 lipidation as post-Golgi vesicles. This is consistent with the sensing of STING-dependent vesicle de-acidification by lyso-pHluorin, a pH sensor usually targeted to endolysosomes. Although it seems likely that STING may neutralize the pH of the Golgi complex, no endogenous LC3 puncta were observed to colocalize with the Golgi markers GM130 or Golgin97. Surprisingly, the presence of C53 blocked STING trafficking at a pre-Golgi stage, leading to STING accumulation in punctate structures that were still capable of activating TBK1 but not LC3 lipidation. The problem can be solved by adding C53 after STING traffics to the Golgi or post-Golgi vesicles, allowing selective inhibition of the ion channel functions of STING by C53.

During the preparation of this manuscript, a similar study by Liu et al. was published^33^. The two studies both uncovered STING-mediated de-acidification of perinuclear vesicles but through different approaches and pH sensors. Both studies observed STING-mediated proton flux in vitro and achieved a block of STING’s ion channel function by C53 that tightly locks the transmembrane pore. Of note, in both studies, purified STING was found to be constitutively active in proton flux, suggesting that STING might be partially activated during purification. Both studies also observed a block of STING-dependent vesicle de-acidification and cell death by C53, while our study established a more strict protocol in assessing the impact of C53 as a STING channel blocker without indirect effects, in addition to identifying post-Golgi vesicles as the major sites of action for the STING channel. Overall, the two studies support the same conclusion that an ion channel function of STING contributes to vesicle pH neutralization, LC3 lipidation, and cell death.

While the cGAS/STING pathway is critical for innate immunity, abnormal activation of this pathway has been increasingly implicated in various human diseases including Parkinson’s and Alzheimer’s diseases, as well as normal aging^34–36^. The ion channel function of STING described here opens an avenue for new therapeutic strategies to target this pathway. Selective manipulation of the inflammation signaling or the ion channel activity depending on the contexts of specific diseases might be considered for optimal therapeutic outcomes.

## Acknowledgements

We thank members of the Shang and Tan laboratories and the Aging Institute members for discussions. We thank Drs. Michael B. Butterworth and Daniel C. Devor from University of Pittsburgh as well as Dr. Qingfeng Chen from Yunnan University for ion channel discussions. We thank Dr. Christian Rosenmund from Charité-Universitätsmedizin Berlin for the lyso-pHluorin plasmid. This work was supported by the National Key R&D Program of China (2021YFC2301400); the National Natural Science Foundation of China (32070876); the Shanxi Provincial Science Fund for Distinguished Young Scholars program (202103021221001) to G.S. This work was also supported by start-up funding from the Aging Institute at the University of Pittsburgh School of Medicine and University of Pittsburgh Medical Center (UPMC) to J.X.T. as well as a UPMC competitive medical research fund award to J.X.T. The authors declare no conflicts of interest.

