## Supplementary information for "A conserved ion channel function of STING mediates non-canonical autophagy and cell death"

### SUPPLEMENTARY MATERIALS

### METHODS

#### Cell culture, chemicals, and treatments

BJ, U2OS, and 293T cells were from ATCC and were authenticated through short tandem repeat (STR) profiling. All relevant authentication data are publicly available from ATCC. These cell lines differ in their growth rates and morphologies. All cell lines in this study were free of contaminations from other cells lines or mycoplasma. Cells were maintained with mycoplasma reagent and potential contaminations are regularly monitored through polymerase chain reaction (PCR) detection. All cells were cultured at 37 °C with 5% $CO_2$ in Dulbecco's modified Eagle's medium (DMEM) supplemented with 8% fetal bovine serum (FBS) and penicillin/streptomycin. Cyclic [G(2',5')pA(3',5')p] or cGAMP (#CT-CGMAP, ChemieTek) was dissolved in water and stored at -20 °C. The other two STING agonists diABZI (#S8796, Selleck) and C53 (#37354, Cayman) were dissolved in DMSO and stored in qliquots at -80 °C. Monensin (#16488, Cayman) was dissolved in ethanol and stored at -20 °C. cGAMP was delivered to cells using a mild digitonin buffer (50 mM HEPES pH 7.0, 100 mM KCl, 3 mM MgCl2, 0.1 mM DTT, 85 mM Sucrose, 0.2% BSA, 1 mM ATP, 8 µg/mL Digitonin)[1]. Cells were treated with the buffer containing 1 µM cGAMP for 5-10 min and then changed back to the original culture media. All other chemicals were directly added to cell culture media.

#### Antibodies

Antibodies for LAMP-1 (sc-20011, IF 1:200); LAMP2 (sc-18822, IF 1:200); Golgin 97 (sc-59820, IF 1: 500); GAPDH (sc-365062, WB 1:5000); Tubulin (sc-5286, WB 1:3000) were from Santa Cruz Biotechnology. The GM130 antibody (610822, IF 1:1000) was from BD Biosciences. Flag (M2, IF 1:1000; F7425, WB 1:3000) antibody was from Sigma. LAMP1 rabbit mAb (#9091, IF 1:200); GM130 rabbit mAb (#12480, IF 1:300); Rab5 rabbit mAb (#3547, IF 1:200), Phospho-STING S366 (#50907, WB 1:1000); Phospho-TBK1 Ser172 rabbit mAb (#5483, WB/IF 1:1000) were from Cell signaling. The following antibodies were from Proteintech: STING (19851-1-AP, WB 1:1000, IF 1:1000); LC3 (18725-1-AP, IF 1:1000, WB 1:1000); TGN46 (13573-1-AP, IF 1:1000); Golgin97 (12640-1-AP, IF 1:1000). Alexa-488/594- and Pacific Blue-conjugated secondary antibodies were obtained from ThermoFisher Scientific.

#### DNA cloning

We used a lentiviral approach for stable protein expression. The DNA sequences of interest were PCR amplified and inserted into a lentiviral vector pCDH-CMV-MCS. When two genes are fused together, a GSGSGS linker was used. Small epitope tags were added directly into PCR primers. To generate point mutations, two fragments of the target cDNA were amplified by PCR with their overlapping ends carrying the intended mutations. The two fragments were then fused together into the pCDH vector through infusion reactions. All new plasmids are verified by DNA sequencing.

#### Stable cell line generation

All experiments in this study are based on stable lines. No transient transfection of DNA was used in any experiment to avoid DNA-induced activation of the cGAS/STING pathway. We used a lentiviral approach for stable protein expression. Lentiviruses were packaged in 293T cells by transfecting the pCDH vector carrying the gene of interest together with the packaging vectors pMD2.G and pSPAX2 using Lipofectamine 2000 (ThermoFisher). The culture media containing the viruses were collected 48 hours after transfection, and were immediately used to infect target cells for stable protein expression. When needed the infected cells were selected by puromycin treatment. The lowest virus titer was used to achieve the desired infection rate.

**Immunofluorescence**

Cells were seeded on glass coverslips (Electron Microscopy Sciences, #7223001) in 24-well plates. After indicated cell stimulations, cells were fixed with 4% Paraformaldehyde (PFA) in *phosphate buffered saline* (PBS) for 30 min at room temperature. Cells were then permeabilized with 0.1% Triton X-100 for 2 min, and blocked for 30 min with 1 x fluorescent blocking buffer (Thermo Fisher Scientific, #37565). The same buffer was further used for the dilution of primary and secondary antibodies in the following cell staining steps. Cells were incubated with primary antibodies at 4 °C overnight or at room temperature for 2 hours. Unbound primary antibodies were washed away with PBS, and cells were further stained fluorescently labeled secondary antibodies. Cells were then washed more than three times to remove any unbound secondary antibodies. When 4′,6-diamidino-2-phenylindole (DAPI) staining was needed, cells were stained with DAPI for 3 min and then washed with PBS. The coverslips were mounted on slides using VECTASHIELD Mounting Medium (Vector Laboratories, #H-1700). Fluorescence images were collected using a Leica SP8 LIGHTNING confocal system with a built-in software Leica Application Suite X 3.5.5.19976. Different positive and negative controls were included to rule out nonspecific staining and any cross talks between channels. Live cell imaging was done using the same confocal system with an Okolabstage-top incubator. All images in the same panel were from the same experiment, followed with the same cell staining, image collection settings, and image processing.

**Immunoblotting**

Cells with indicated treatments were briefly washed with cold PBS and then lysed with a lysis buffer containing 50 mM Tris-HCl, pH7.5, 150 mM NaCl, 0.5% Triton X-100, 2 mM NaF, and protease inhibitor cocktail. The lysates were briefly sonicated to fully dissolve all membranes, followed by centrifugation at 15,000 xg for 10 min. The supernatants were collected and heated at 95 °C for 5 min after mixing with equal volumes of 2x SDS loading buffer (0.1 M Tris HCl, pH 6.8, 4% SDS, 20% glycerol, 2% 2-Mercaptoethanol, 0.01% bromphenol blue). The protein samples were kept at -80 °C or directly moved to immunoblotting analysis. Protein electrophoresis was done using 4-20% precast polyacrylamide gel (Biorad, #4561096), followed by protein transfer to 0.45 µm Nitrocellulose membranes using the Trans-Blot Turbo system. The membranes were then blocked with StartingBlock Blocking Buffer (ThermoFisher, #37542) for 30 min and then incubated sequentially with primary and secondary antibodies. After washing, the target proteins were detected Immobilon Forte Western HRP Substrate (Sigma-Aldrich, # WBLUF0500) in a ChemiDoc MP Imaging System.

**Lyso-pHluorin assay**

Cells stably expressing Lyso-pHluorin at 50-70% confluency were treated with vehicle or indicated STING agonists to trigger STING trafficking and vesicle de-acidification. For cGAMP treatment, the media from the cell culture dish were moved to a new tube and kept at 37 °C. Cells were treated with the digitonin/cGAMP buffer for 10 min, and the buffer was then replaced with the original warm culture media. Other chemicals were directly added to and mixed well in culture media. Cells were subsequently monitored for changes in their lyso-pHluorin signals every 30 min and live cell images were collected at desired time points.

**Protein expression and purification**

The human STING protein was expressed and purified following established protocols[2,3]. Briefly, The coding sequences of human STING gene was inserted into modified pEZT-BM vector, generating a STING fusion protein with C-terminal Tsi3 tag[4]. The plasmid was transfected into HEK293F cells using PEI with a mass ratio of 1 : 3 (plasmid : PEI) and the expression was enhanced by 3 mM sodium butyrate 12 h later. After 48 h of culturing, cells were collected and re-suspended into buffer A (20 mM Tris pH 7.5, 150 mM NaCl, 1 mM AEBSF and protease inhibitor cocktail). Then, cells were lysed by French press and the membranes were collected using ultracentrifugation at 10,0000 g. The membrane proteins were extracted using 1% DDM/CHS (5:1) and the insoluble part was removed by ultracentrifugation at 10,0000 g. The supernatant was loaded onto the Tse3-conjugated resin pre-equilibrated with buffer B (20 mM Tris pH 8.0, 150 mM NaCl, 1mM $CaCl_2$, 0.03% DDM and

0.003% CHS). The Tsi3 tag was removed using on-column digestion by adding 3C-protease overnight. The STING protein was further polished by applying to size exclusion chromatography (SEC) in buffer C (20 mM HEPES pH 7.5, 150 mM NaCl, 0.03% DDM, 0.003% CHS and 1 mM TCEP).The peak was pooled, concentrated at 8 mg/ml and stored at -80 °C for flux assay.

**Fluorescence-based proton flux assay.**

Lipids of PC (20 mg/ml), PE (20 mg/ml) and PE (20 mg/ml) were mixed in chloroform at a 2:1:1 ratio and dried using a nitrogen stream and trace mount of chloroform was removed using a vacuum chamber. Dried lipids were suspended in internal buffer C (50 mM HEPES pH 7.4, 450 mM KCl and 5 mM DTT) generating the liposome with a concentration of 10 mg/ml. The unilamellar liposomes was formed by freeze-thawing cycle followed by sonication. An aliquot of liposomes were mixed with 0.7% DDM for 2 h at 25 °C. Human STING protein was then added to the liposome/DDM mixture (ratio 1: 100) and incubated for 1 h at 25 °C. DDM was completely removed by adding Bio-beads SM2 (Bio-rad) every 4 h at 4 °C for 3 times. Generated proteoliposome vesicles were collected and used for ion channel flux assays. Proteoliposomes (12 µl) were mixed with 160 µl external flux assay buffer D (5 mM HEPES pH 7.4, 450 mM NaCl, 5 mM DTT and 2 uM ACMA) in a 96-well fluorescence assay plate. AMCA fluorescence intensity was measured over time ($\lambda_{Ex}$ = 410 nm, $\lambda_{Em}$ = 490 nm). The 0.45 uM valinomycin was added to initiate the flux and fluorescence data were collected at 20-second intervals for 15 min and then 1 µM CCCP was used to collapse the proton gradient. Data were processed according to established protocol[5].

**Image analysis**

For the quantification of fluorescence images, the outlines of randomly selected cells were manually annotated in each image by an investigator blinded to the allocation. The fraction of protein A's intensity on protein B was quantified by comparing the sum of signal intensities of protein A overlapping with protein B to the total intensity of protein A in a given cell. When quantifying the fraction of protein A positive for protein B puncta, a higher threshold was applied to minimize diffuse protein B signals. Thresholds for both proteins were determined by the target protein's signal intensity percentile within a single cell, adjusted by a small constant. All cells from the same experiments were applied using the same threshold settings. Manual checking ensured accurate threshold applications. Quantitative data were exported to Excel and visualized using Prism to display mean and SEM values.

**Software**

A Leica SP8 LIGHTNING confocal system and its built-in software Leica Application Suite X 3.5.5.19976 were used to collect confocal images as well as for living cell imaging. Images were processed in Adobe Photoshop 20.0.4. Schematic illustration figures were made with Biorender and Microsoft Office Power Point Professional Plus 2016. Protein structures were visualized in the PyMOL, Version 2.4.0. Graphs were generated in GraphPd Prism 9.0.0. Images were quantified using custom-coded algorithms tailored according to specific experiments to incorporate quality control measures, minimize errors, and ensure accuracy and reliability.

**Statistics and reproducibility**

All experiments were independently reproduced at least three times unless otherwise indicated. Strict standards were applied to screen for robust and unbiased results. No statistical methods were used to predetermine sample size. The investigators were blinded to allocation during data collection and image quantification. Data were presented as mean ± SEM. Statistical significance was determined by unpaired, two-tailed t-tests.

**Data Availability**

All data of this study are included in this paper. Additional information is available upon request. Custom codes for image quantifications are available at https://github.com/jaytanlab/Protein_Colocalization_Quantification.

### EXTENDED DATA FIGURE LEGENDS

**Extended Data Figure 1**. **STING traffics through the Golgi, causing morphological changes of the Golgi stacks.**

**A.** Compound C53 inhibits diABZI-induced LC3 lipidation in BJ cells without blocking TBK1 signaling and STING phosphorylation. BJ cells treated as indicated were harvested 3 hours after treatment for western blot.

**B.** STING trafficking triggers morphological changes of the Golgi stacks 30 min after cGAMP stimulation in U2OS cells. Note brighter and swelled TGN marker Golgin97 at 30 min. Monoclonal U2OS Flag-hSTING cells were stimulated with 1 μM cGAMP and fixed at indicated time points for the co-staining of endogenous *cis*- and *trans*-Golgi markers GM130 and Golgin97, respectively.

**C, D.** Quantification of the colocalization between GM130 and Golgin97 in (B). Mean ± SEM; n = 68, 96, 50, and 86 random cells for 0', 30', 60', and 90', respectively.

**E.** STING trafficking triggers morphological changes of the Golgi stacks 30 min after cGAMP stimulation in HT1080 cells expressing endogenous STING. Note brighter and swelled TGN marker Golgin97 as well as increased overlap between GM130 and Golgin97 at 30 min. HT1080 cells were stimulated with 1 μM cGAMP and fixed at indicated time points for the co-staining of endogenous *cis*- and *trans*-Golgi markers GM130 and Golgin97, respectively.

**F.** STING trafficking causes TGN46 budding onto Gogli-derived vesicles which partially colocalize with post-Golgi STING vesicles. Monoclonal U2OS Flag-hSTING cells were stimulated with 1 μM cGAMP and fixed at indicated time points for the co-staining of STING and TGN46. Note, TGN46 was found out of the Golgi stacks at 30 min before STING budded out from TGN.

**G.** diABZI stimulates STING trafficking through the Golgi in U2OS cells. Monoclonal U2OS Flag-hSTING cells were stimulated with 1 μM diABZI and fixed at indicated time points for the co-staining of STING and Golgin97.

**H.** Quantification of the colocalization between STING and Golgin97 in (G). Mean ± SEM; n = 57, 65, 76, 74, and 66 random cells for 0', 30', 60', 120', and 180', respectively.

Bar, 10 μm.

**Extended Data Figure 2**. **Human and mouse STING similarly traffic through the Golgi complex.**

**A, B.** STING traffics through the Golgi after cGAMP stimulation in HT1080 (A) and BJ (B) cells, both expressing endogenous human STING. Cells were stimulated with 1 μM cGAMP and fixed at indicated time points for the co-staining of endogenous STING and the *trans*-Golgi marker Golgin97, respectively. Note, brighter and swelled TGN marker Golgin97 at 30 min in HT1080 cells, whereas STING accumulation at the Golgi in BJ cells appeared to peak between 30 and 60 min.

**C.** Mouse STING traffics through the *trans*-Golgi upon cGAMP binding. U2OS cells stably expressing low levels of Flag-mSTING were stimulated with 1 μM cGAMP and fixed at indicated time points for the co-staining of STING and the *trans*-Golgi marker Golgin97.

**D, E.** Quantification of the colocalization between STING and Golgin97 in (C). Mean ± SEM; n = 49, 48, 36, and 52 random cells for 0', 30', 60', and 90', respectively.

**F.** STING traffics through the *cis*-Golgi upon cGAMP binding. U2OS cells stably expressing low levels of Flag-mSTING were stimulated with 1 μM cGAMP and fixed at indicated time points for the co-staining of STING and the cis-Golgi marker GM130.

**G, H.** Quantification of the colocalization between STING and GM130 in (F). Mean ± SEM; n = 41, 50, 51, and 59 random cells for 0’, 30’, 60’, and 90’, respectively.

Bar, 10 μm.

**Extended Data Figure 3**. **Post-Golgi STING vesicles develop endosome-like properties accompanied by LC3 lipidation.**

**A.** STING induces LC3 puncta outside of the Golgi. Monoclonal U2OS Flag-hSTING cells were stimulated with 1 μM cGAMP and fixed at indicated time points for the co-staining of endogenous LC3B and the *cis*-Golgi marker GM130.

**B.** Quantification of the colocalization between LC3 and GM130. Mean ± SEM; n = 73, 79, 75, 86, 40, 55, 87, and 71 random cells from left to right.

**C.** STING induces LC3 puncta outside of the Golgi in BJ cells. Cells were stimulated with 1 μM cGAMP and fixed at indicated time points for the co-staining of endogenous LC3B and the *trans*-Golgi marker Golgin97.

**D.** LC3 puncta were found on STING vesicles leaving the perinuclear vesicle clusters and were not found on Golgin97. BJ cells stably expressing EGFP-LC3B were fixed 90 min after cGAMP stimulation for immunostaining of STING and Golgin97.

**E.** EGFP-LC3B colocalizes with STING but not Golgin97. Normalized fluorescence intensities of Golgin97, EGFP-LC3B, and STING along the white line in the right bottom image of panel (D).

**F.** LC3 puncta were found on STING vesicles positive for RAB5, near the periphery of the post-Golgi vesicle clusters. BJ cells stably expressing EGFP-LC3B were fixed 90 min after cGAMP stimulation for immunostaining of STING and RAB5.

**G.** STING puncta develop a relatively low level of colocalization with the late endosome/lysosome marker CD63. BJ Cells were stimulated with 1 μM cGAMP and fixed at indicated time points for the co-staining of endogenous STING and CD63.

**H, I.** Quantification of the colocalization between STING and CD63 in (G). Mean ± SEM; n = 16, 16, 20, 23, and 28 random cells from left to right.

**J.** STING puncta develop a relatively low level of colocalization with the late endosome/lysosome marker LAMP1. BJ Cells were stimulated with 1 μM cGAMP and fixed at indicated time points for the co-staining of endogenous STING and CD63.

**K, L.** Quantification of the colocalization between STING and LAMP1 in (J). Mean ± SEM; n = 27, 23, 37, 29, 24, and 24 random cells from left to right.

Bar, 10 μm.

**Extended Data Figure 4**. **STING neutralizes the pH of post-Golgi vesicles, which is captured lyso-pHluorin, an endolysosomal pH sensor.**

**A.** cGAMP stimulates lyso-pHluorin puncta in wild type BJ cells. BJ cells stably expressing lyso-pHluorin were treated with digitonin buffer alone (Vehicle) or with cGAMP for 10 min, changed back to original media, and chased for indicated time periods. Lyso-pHluorin puncta were monitored by live cell imaging.

**B.** STING activation does not induce EGFP-galectin3 puncta. U2OS cells stably expressing EGFP-galectin3 and human or mouse STING (hSTING/mSTING) were stimulated with indicated STING agonists. The fluorescence of EGFP-galectin3 were monitored by live cell imaging.

**C.** Setting up a protocol to fix lyso-pHluorin cells for the immunostaining of other proteins without triggering new lyso-pHluorin puncta by fixation or permeabilization. Monensin was used to induce pre-existing puncta before fixation. 30 min of fixation in 4% electron microscopy grade polyformaldehyde (PFA) followed by 2 min of permeabilization in 0.1% Triton-X 100 in PBS were used as a standard protocol to examine the colocalization of lyso-pHluorin puncta with other proteins by immunostaining.

**D, E.** Normal Lyso-pHluorin puncta formation upon mSTING activation by DMXAA in ATG5-KO (D) and ATG7-KO (E) U2OS cells stably expressing mSTING and lyso-pHluorin.

Bar, 10 µm.

**Extended Data Figure 5**. **Testing STING channel mutants.**

**A.** The side view of human STING (PDB: 8IK3) with the channel entry at the cytosolic side. The electronic potential was shown. Blue and red colors represent the positively and negatively charged regions, respectively.

**B.** Bottom view of cGAMP bound human STING (PDB: 8IK3). The protomers are shown in cartoon with yellow and blue color, respectively. The residues lining up the pore are labelled.

**C.** The list of STING pore mutations tested in this study.

**D.** No basal lyso-pHluorin puncta were observed in U2OS cells infected by lentiviruses to stably express each STING mutants.

**E.** Most pore mutants of STING still triggered lyso-pHluorin puncta after cGAMP stimulation except Mutants #8 and #18 which were not expressed as shown in panel (F). Note that Mutant #3 (L54E) induced brighter puncta than wild-type or other mutants of STING.

**F.** Western blot analysis of the protein levels of each pore mutants of STING in U2OS cells.

**G.** The localization of L54 around the pore in the structure of cGAMP-bound human STING (PDB: 8IK3). The pore radii (spheres) were calculated using HOLE.

Bar, 10 µm.

**Extended Data Figure 6**. **Compound C53 can be used as a STING channel blocker when used after STING traffics to the Golgi or post-Golgi vesicles.**

**A.** Compound C53, which binds to the transmembrane pore of STING, induces STING puncta that do not colocalize with the Golgi. Monoclonal U2OS Flag-hSTING cells were stimulated with C53 alone and then fixed at indicated time points for immunostaining of STING and the trans-Golgi marker Golin97.

**B.** Compound C53 fully blocks cGAMP-induced trafficking of Flag-STING from the ER to the Golgi in U2OS cells. Monoclonal U2OS Flag-hSTING cells were stimulated with cGAMP + C53 for indicated time periods, followed by fixation and immunostaining of STING and the cis-Golgi marker GM130. See quantification in (B).

**C.** Quantification of the colocalization between STING and GM130. Mean ± SEM; n = 49, 58, 109, 147, 58, 27, 115, 96, and 93 random cells from left to right.

**D.** C53 fully blocks diABZI-induced trafficking of Flag-STING from the ER to the Golgi in U2OS cells. Monoclonal U2OS Flag-hSTING cells were stimulated with diABZI + C53 for indicated time periods, followed by fixation and immunostaining of STING and the trans-Golgi marker Golgin97.

**E.** C53 addition 30 min after cGAMP allows STING trafficking to the Golgi and post-Golgi vesicles. Monoclonal U2OS cells stably expressing hSTING were stimulated as indicated and then fixed for the immunostaining of STING and GM130. See quantification in (B).

**F.** C53 addition 1 hour after diABZI fully blocks STING-dependent LC3 lipidation. Monoclonal U2OS Flag-hSTING cells were stimulated as indicated and whole cell lysates were harvested for western blot 2.5 hours after diABZI treatment.

**G.** C53 addition 2 hours after cGAMP stimulation failed to block STING-dependent LC3 lipidation. The same cells in (E) were treated as indicated and harvested for western blot.

Bar, 10 μm.

Fig S1

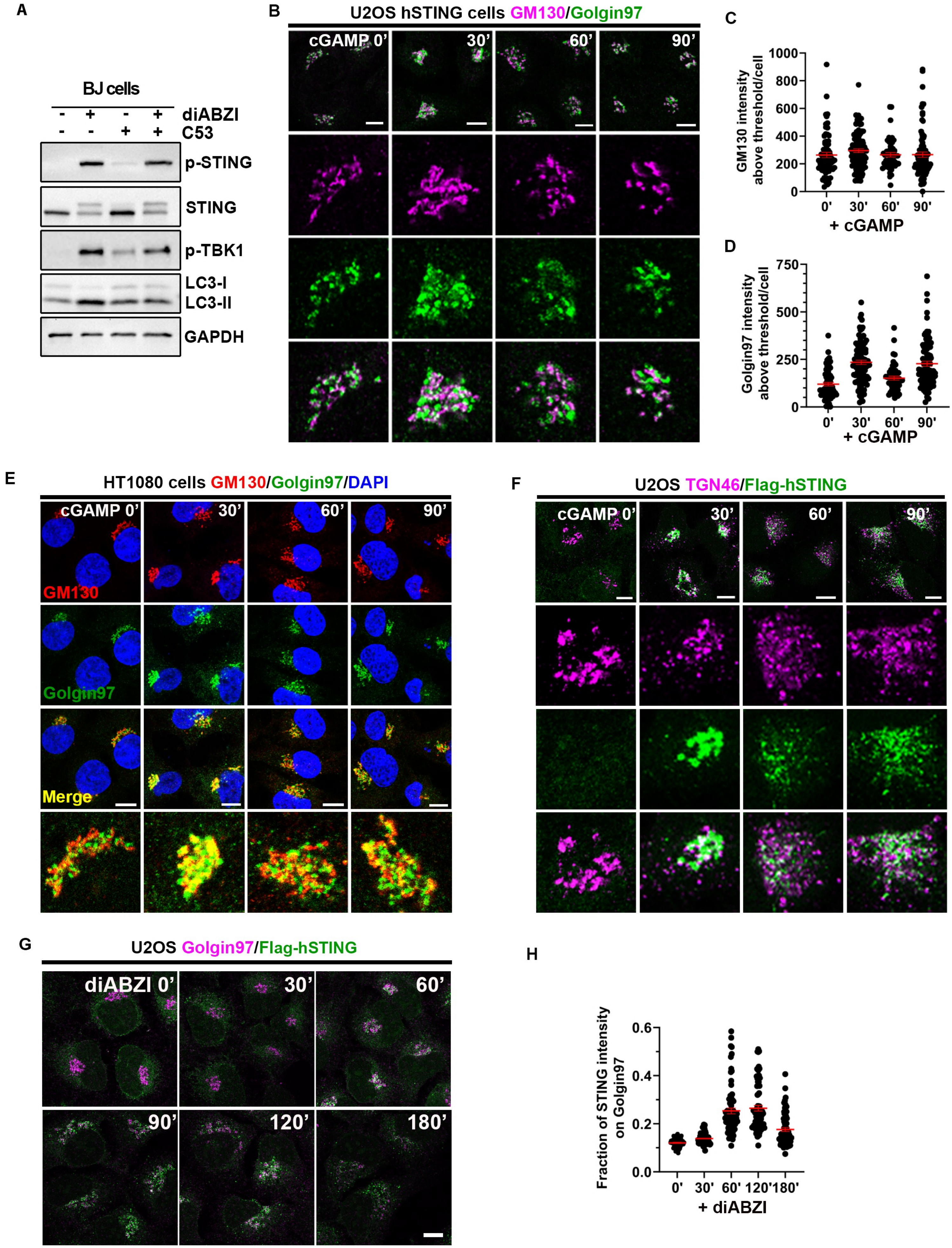

A
BJ cells
diABZI
C53
p-STING
STING
p-TBK1
LC3-I
LC3-II
GAPDH
B
U2OS hSTING cells GM130/Golgin97
cGAMP 0'
30'
60'
90'
C
GM130 intensity above threshold/cell
1000
800
600
400
200
0
0'
30'
60'
90'
+ cGAMP
D
Golgin97 intensity above threshold/cell
750
500
250
0
0'
30'
60'
90'
+ cGAMP
E
HT1080 cells GM130/Golgin97/DAPI
cGAMP 0'
30'
60'
90'
GM130
Golgin97
Merge
F
U2OS TGN46/Flag-hSTING
cGAMP 0'
30'
60'
90'
G
U2OS Golgin97/Flag-hSTING
diABZI 0'
30'
60'
90'
120'
180'
H
Fraction of STING intensity on Golgin97
0.6
0.4
0.2
0.0
0'
30'
60'
120'
180'
+ diABZI

Fig S2

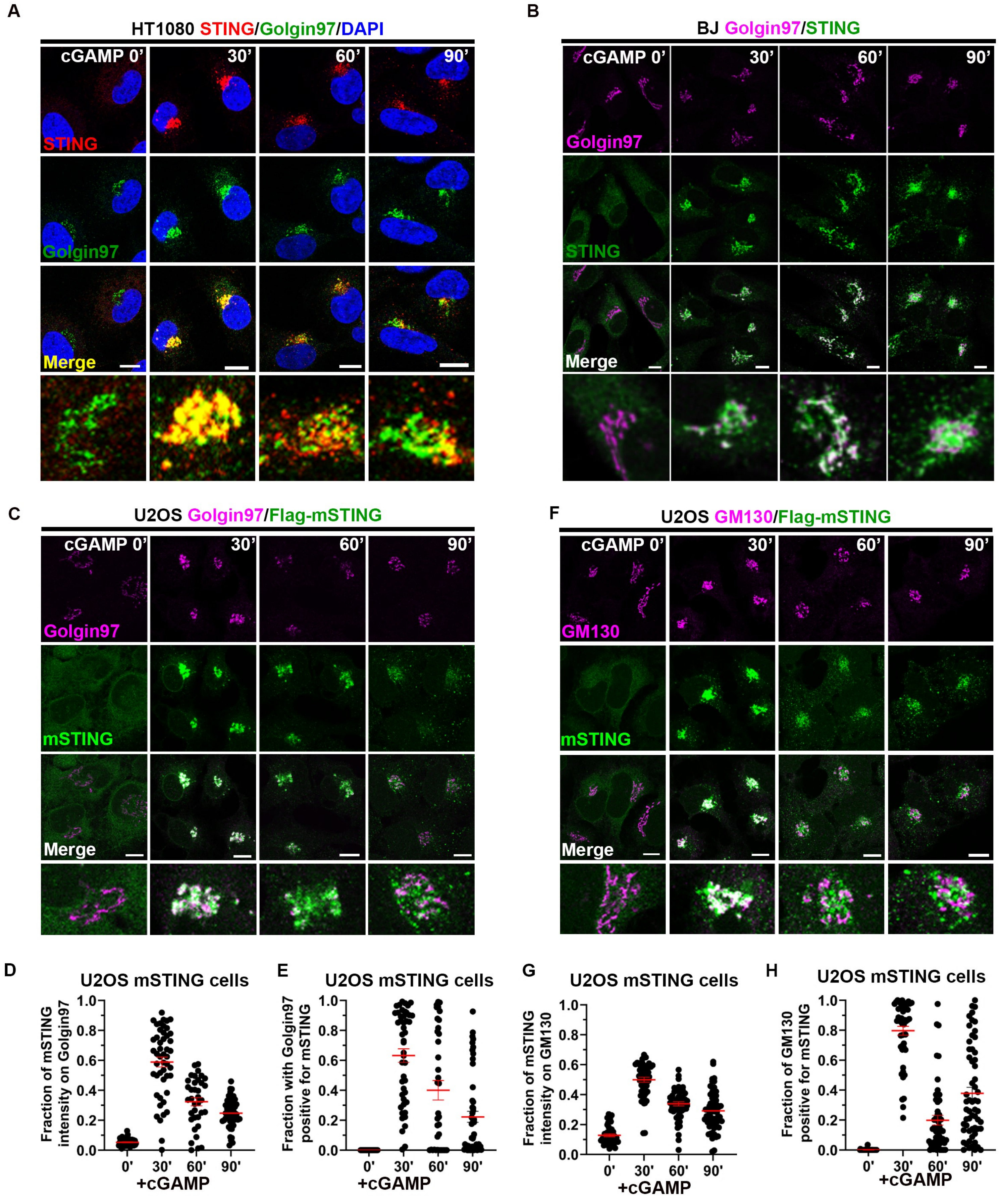
A
HT1080 STING/Golgin97/DAPI
cGAMP 0'
30'
60'
90'
STING
Golgin97
Merge
B
BJ Golgin97/STING
Golgin97
STING
Merge
C
U2OS Golgin97/Flag-mSTING
Golgin97
mSTING
Merge
F
U2OS GM130/Flag-mSTING
GM130
mSTING
Merge
D
U2OS mSTING cells
Fraction of mSTING intensity on Golgin97
+cGAMP
E
U2OS mSTING cells
Fraction with Golgin97 positive for mSTING
+cGAMP
G
U2OS mSTING cells
Fraction of mSTING intensity on GM130
+cGAMP
H
U2OS mSTING cells
Fraction of GM130 positive for mSTING
+cGAMP
0.0
0.2
0.4
0.6
0.8
1.0
0'
30'
60'
90'

Fig S3

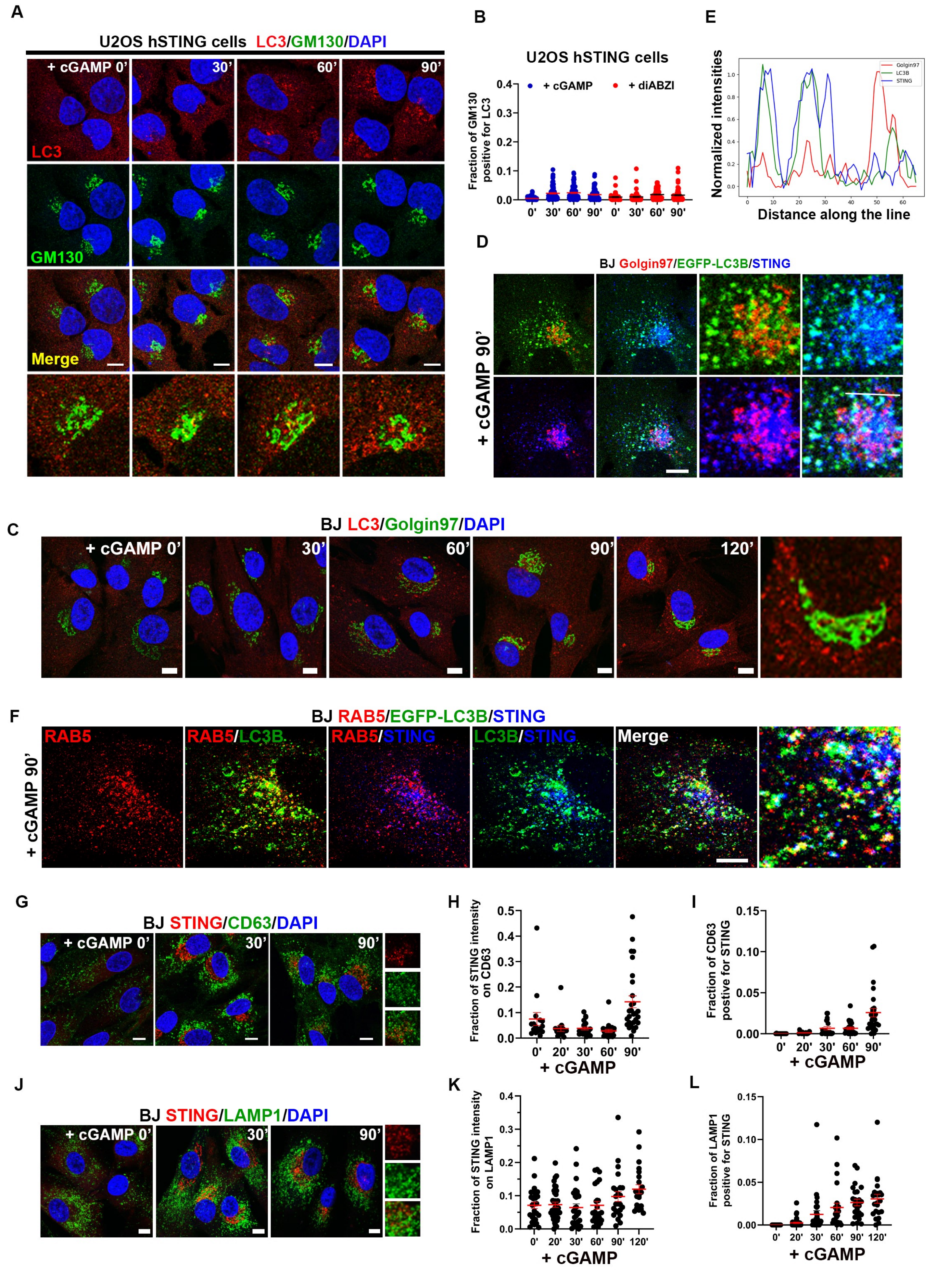
A
U2OS hSTING cells LC3/GM130/DAPI
+ cGAMP 0'
30'
60'
90'
LC3
GM130
Merge
B
U2OS hSTING cells
+ cGAMP
+ diABZI
Fraction of GM130 positive for LC3
0'
30'
60'
90'
E
Normalized intensities
Golgin97
LC3B
STING
Distance along the line
D
BJ Golgin97/EGFP-LC3B/STING
+ cGAMP 90'
C
BJ LC3/Golgin97/DAPI
+ cGAMP 0'
30'
60'
90'
120'
F
BJ RAB5/EGFP-LC3B/STING
RAB5
RAB5/LC3B
RAB5/STING
LC3B/STING
Merge
+ cGAMP 90'
G
BJ STING/CD63/DAPI
+ cGAMP 0'
30'
90'
H
Fraction of STING intensity on CD63
0'
20'
30'
60'
90'
+ cGAMP
I
Fraction of CD63 postive for STING
0'
20'
30'
60'
90'
+ cGAMP
J
BJ STING/LAMP1/DAPI
+ cGAMP 0'
30'
90'
K
Fraction of STING intensity on LAMP1
0'
20'
30'
60'
90'
120'
+ cGAMP
L
Fraction of LAMP1 positive for STING
0'
20'
30'
60'
90'
120'
+ cGAMP

Fig S4

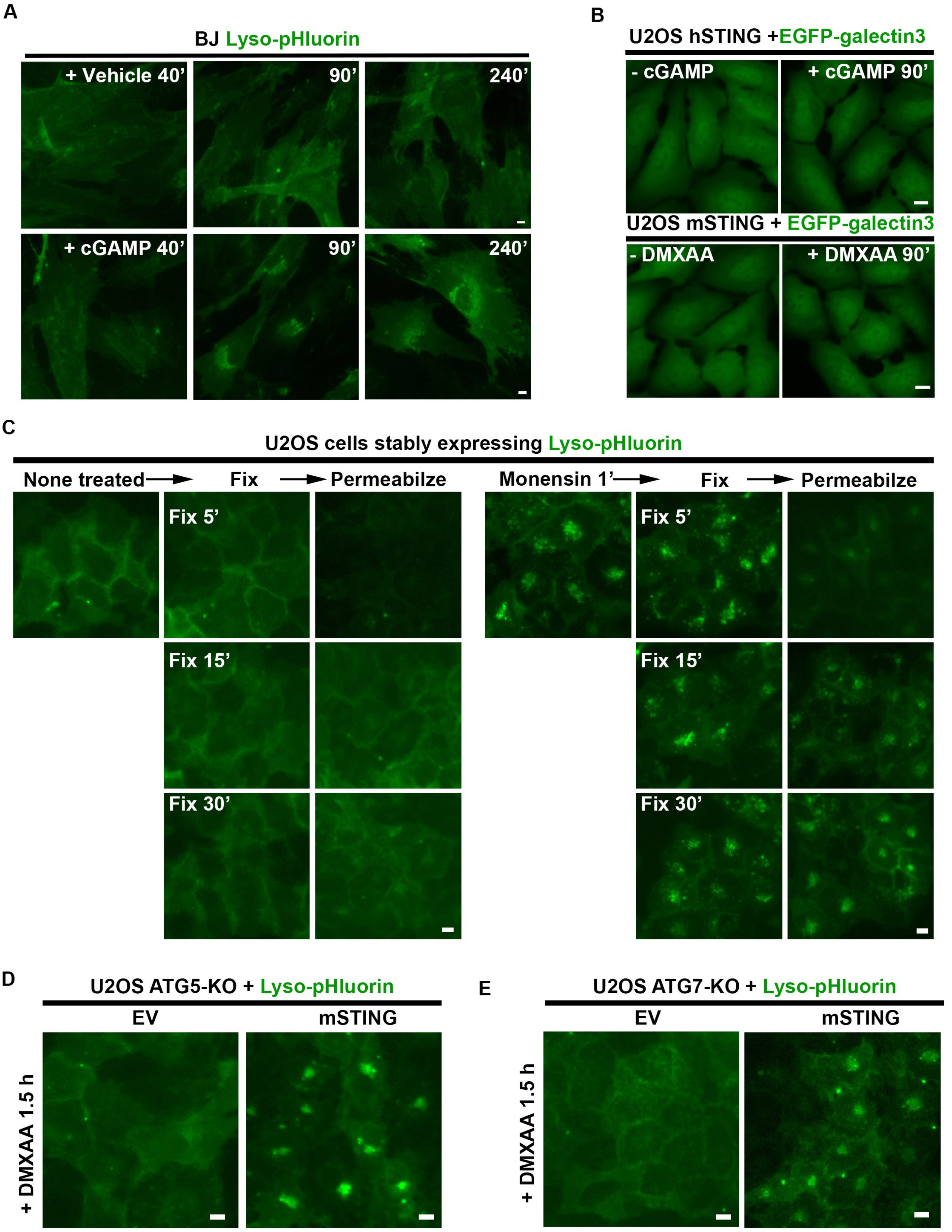
A
BJ Lyso-pHluorin
+ Vehicle 40'
90'
240'
+ cGAMP 40'
90'
240'
B
U2OS hSTING +EGFP-galectin3
- cGAMP
+ cGAMP 90'
U2OS mSTING + EGFP-galectin3
- DMXAA
+ DMXAA 90'
C
U2OS cells stably expressing Lyso-pHluorin
None treated
Fix
Permeabilze
Monensin 1'
Fix
Permeabilze
Fix 5'
Fix 15'
Fix 30'
Fix 5'
Fix 15'
Fix 30'
D
U2OS ATG5-KO + Lyso-pHluorin
EV
mSTING
+ DMXAA 1.5 h
E
U2OS ATG7-KO + Lyso-pHluorin
EV
mSTING
+ DMXAA 1.5 h

**Fig. S5**

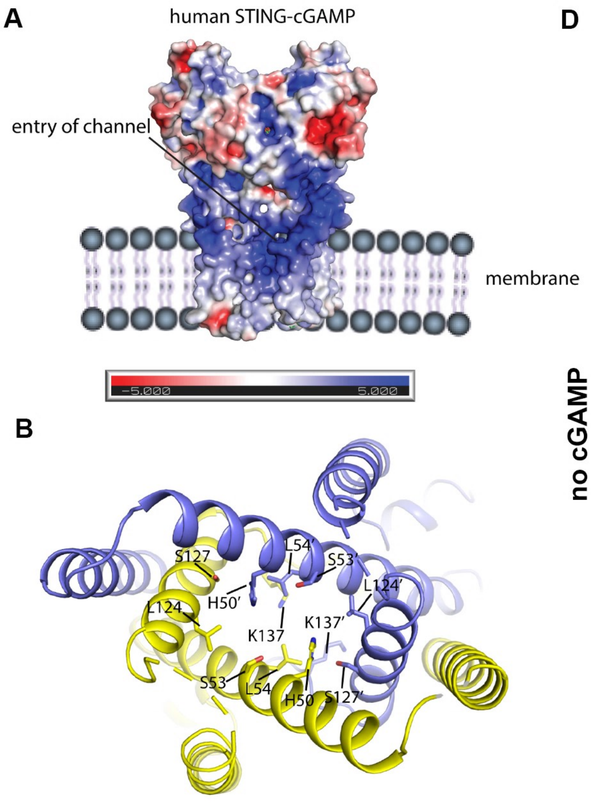


C

| | STING pore mutations |
|---|---|
| 1 | H50L |
| 2 | S53L |
| 3 | L54E |
| 4 | K137E |
| 5 | H50A |
| 6 | H50E |
| 7 | H50Q |
| 8 | H50R |
| 9 | H50W |
| 10 | H50Y |
| 11 | S53M |
| 12 | S53W |
| 13 | S53Y |
| 14 | L54W |
| 15 | H50W S53W |
| 16 | H50Y S53Y |
| 17 | H50W/S53W/L124F |
| 18 | H50W/S53W/S127L |

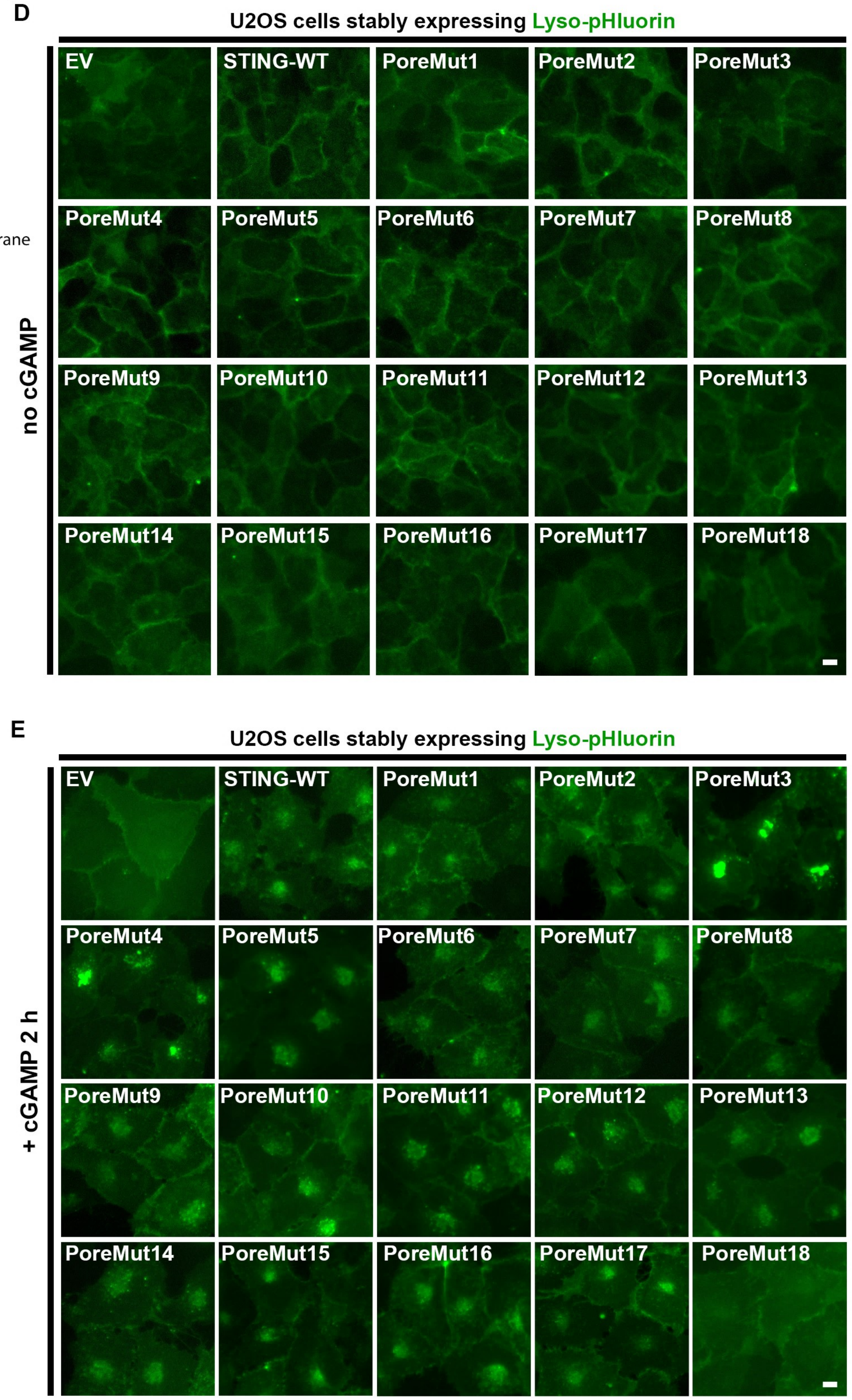


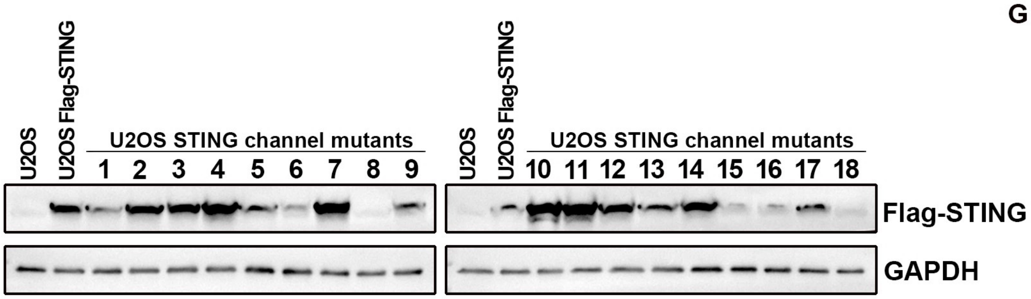


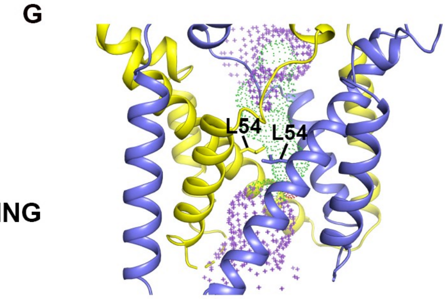

Fig S6

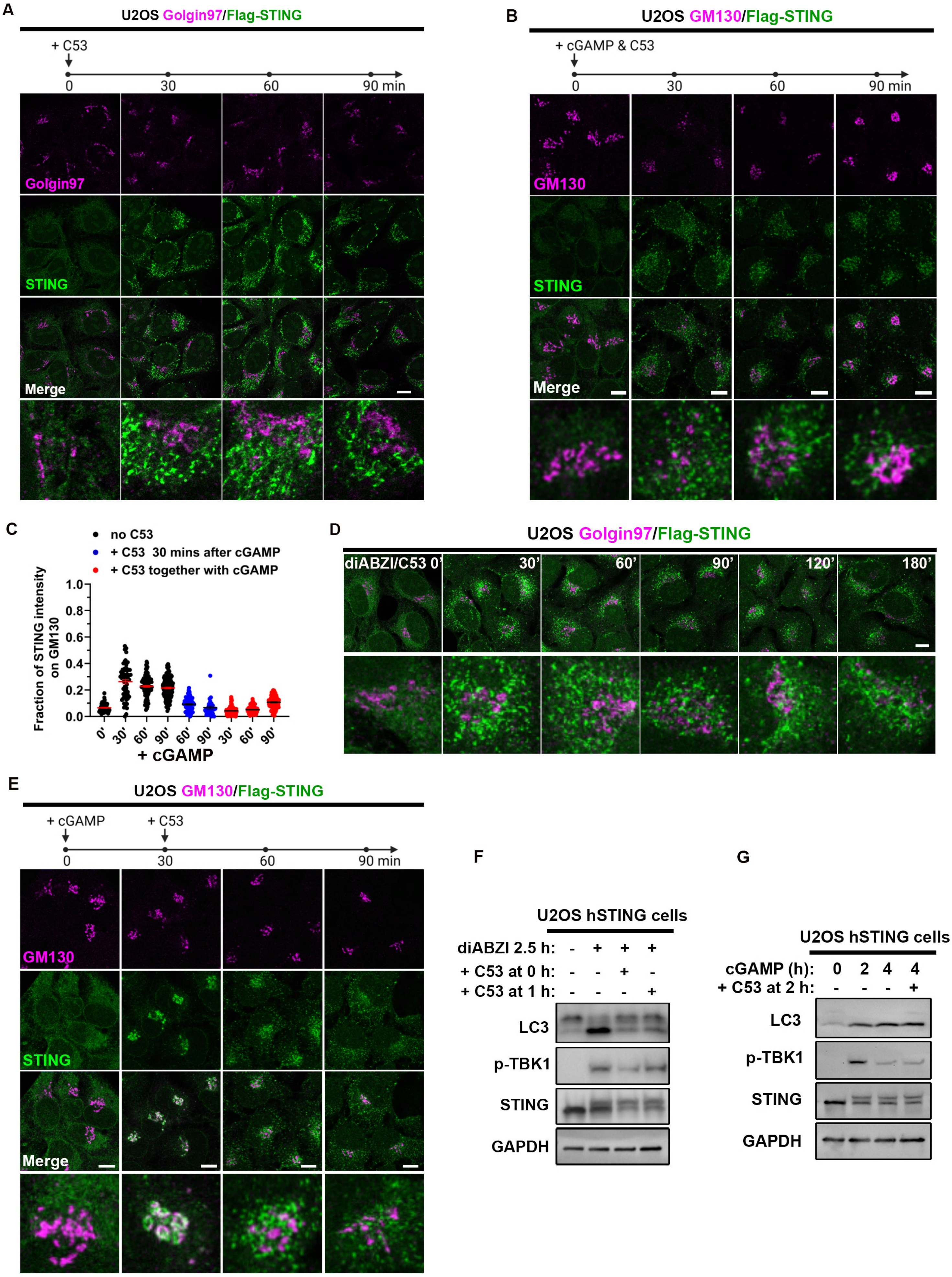
A
U2OS Golgin97/Flag-STING
+ C53
0
30
60
90 min
Golgin97
STING
Merge
B
U2OS GM130/Flag-STING
+ cGAMP & C53
0
30
60
90 min
GM130
STING
Merge
C
no C53
+ C53 30 mins after cGAMP
+ C53 together with cGAMP
Fraction of STING intensity on GM130
1.0
0.8
0.6
0.4
0.2
0.0
0'
30'
60'
90'
60'
90'
30'
60'
90'
+ cGAMP
D
U2OS Golgin97/Flag-STING
diABZI/C53 0'
30'
60'
90'
120'
180'
E
U2OS GM130/Flag-STING
+ cGAMP
+ C53
0
30
60
90 min
GM130
STING
Merge
F
U2OS hSTING cells
diABZI 2.5 h: - + + +
+ C53 at 0 h: - - + -
+ C53 at 1 h: - - - +
LC3
p-TBK1
STING
GAPDH
G
U2OS hSTING cells
cGAMP (h): 0 2 4 4
+ C53 at 2 h: - - - +
LC3
p-TBK1
STING
GAPDH
